# Enhancing Supercooled Red Blood Cell Storage: The Membrane-Stabilizing Effect of Ethanol

**DOI:** 10.64898/2026.07.29.741492

**Authors:** Nishaka William, Ziya Isiksacan, Travis Nemkov, Luke E. Boudreau, Matilda Holtz, Yuanheng Zhao, Jayme Kurach, Mahsa Yazdanbakhsh, Jason P. Acker, Angelo D’Alessandro, O. Berk Usta

**Affiliations:** Laboratory Medicine and Pathology, University of Alberta, Edmonton, AB T6G 2R8, Canada; Center for Engineering in Medicine and Surgery, Department of Surgery, Massachusetts General Hospital, Harvard Medical School, Boston, MA, 02114, USA; Shriners Children’s Hospital, Boston, MA, 02114, USA; Department of Biochemistry and Molecular Genetics, University of Colorado Anschutz Medical Campus, Aurora, CO, 80045, USA; Innovation and Portfolio Management, Canadian Blood Services, Edmonton, AB T6G 2R8, Canada

**Keywords:** Red blood cell, Supercooling, Lipidomics, Metabolomics, Ethanol, Membrane Fluidity, Biopreservation

## Abstract

Hypothermic storage is constrained by the progressive depletion of energy reserves, curtailing the shelf-life of organs, cell therapies, and blood products. High sub-zero *‘supercooling’* helps preserve energy homeostasis by slowing catabolic processes; however, the resulting injury in this setting is not primarily driven by energy depletion. Here, we investigated whether low-dose ethanol could prevent forms of injury that arise independently of disrupted energy homeostasis and remain unaddressed in supercooled storage. Human red blood cells treated with 4% (v/v) ethanol were stored 4 °C, −4 °C, or −8 °C and subject to a series of functional assessments and integrated metabolomic/lipidomic profiling after 21 and 42 days of storage. Metabolomics data showed that energy homeostasis was better preserved at lower temperatures, yet these supercooled conditions simultaneously intensified hemolysis and caused a marked depletion of lysophospholipid species that did not occur at 4 °C. Ethanol blunted these effects, cutting hemolysis by ∼50 % at −4 °C, by ∼85 % at −8 °C, and attenuating lysophospholipid depletion. These results uncover a previously unrecognized, lipid-centric injury that arises during supercooled storage and establish low-dose ethanol as a simple, readily deployable countermeasure that could help extend storage intervals of diverse biological systems.

**SUMMARY:** Cellular preservation has traditionally focused on maintaining energy metabolism during hypothermic storage, but whether this is sufficient at high sub-zero temperatures remains unclear. Human red blood cells were stored for up to 42 days at 4°C, −4°C, or −8°C with or without 4% ethanol and evaluated using functional assays and integrated metabolomic and lipidomic profiling. Although supercooling preserved energy homeostasis, it increased hemolysis and caused pronounced lysophospholipid depletion. Ethanol reduced hemolysis by approximately 50% at −4°C and 85% at −8°C, attenuated lysophospholipid loss, and produced comparatively modest changes in cellular metabolism. These findings identify membrane integrity as a critical determinant of preservation outcome and support strategies that protect membrane stability alongside metabolic homeostasis.

## INTRODUCTION

The deterioration in quality observed during the storage of clinically relevant cell or tissue systems at hypothermic temperatures (1 °C–6 °C) stems from the depletion of energy reserves promoting ion and redox imbalances that, when excessive, cause irreversible cell damage which trigger apoptotic or necrotic pathways ^1^. Hypothermic storage solutions are carefully formulated to slow the depletion of energy reserves while enhancing cellular tolerance to inevitable ion and redox imbalances ^2^. Yet the protective capacity of these solutions is intrinsically constrained; because hypothermia can only slow biological reaction kinetics to a finite degree, surpassing this barrier remains challenging. Advancing biological storage therefore requires new paradigms, with *supercooling* at high sub-zero temperatures emerging as a promising approach which slows catabolic processes which disrupt cellular homeostasis during hypothermic storage and eliminating the need to guard against ice-induced injury.

While supercooling has been shown to extend storage duration in various cell and organ systems, its homeostatic effects are not exclusively beneficial due to increased metabolic uncoupling compared to hypothermic storage ^3–5^. During supercooling, the disparities in reaction rates become more pronounced because the sensitivity of each reaction to temperature hinges on both its thermodynamic drive (enthalpy- vs. entropy-dominated) and the magnitude of its kinetic activation barrier ^6,7^. In practice, this can be harmful: spontaneous, low-barrier modifications – such as protein oxidation, glycation, or deamidation – remain comparatively active at low temperatures, whereas essential repair pathways that rely on larger entropic contributions and higher activation energies slow disproportionately, allowing irreversible damage to accumulate ^6,7^. Compounding this imbalance, supercooling further restricts lateral diffusion in the lipid bilayer, disrupting protein–protein and protein–lipid interactions and disproportionately inactivating membrane-bound enzymes beyond classical Arrhenius predictions ^8^. Fully harnessing supercooled preservation requires curbing this amplified metabolic uncoupling to benefit from conserved energy reserves without impairing protective cellular mechanisms. It is possible that short-chain alcohols could help to meet this need. While traditional cryoprotectants like glycerol and DMSO prevent ice-induced injury during deep freezing, they are sub-optimal for supercooling due to increased bulk viscosity and poor membrane intercalation ^9,10^. Conversely, short-chain alcohols possess the amphiphilicity required to enhance low-temperature membrane fluidity, and early studies have hinted at their superior performance in high-sub-zero storage ^4,11^.

Because RBCs lack extensive *de novo* synthesis and turnover mechanisms, their minimalist biology offers exceptional clarity into the sequelae of storage. Under exogenous stress, RBCs undergo a progressive decline in function, and their metabolic profile tracks this deterioration more faithfully than that of nucleated cells ^12–14^. Our earlier work showed that supercooling (at −4 °C to −8 °C) preserves adenosine triphosphate (ATP) compared with hypothermic storage yet accelerates membrane damage and subsequent hemolysis over the course of six weeks ^15,16^. Employing standard countermeasures that have been used to mitigate injury during hypothermic storage (e.g., antioxidants, solutions with increased pH and reduced osmolarity) only partially attenuate the storage lesion during supercooling; they neither equalize, nor surpass the outcomes observed under conventional hypothermic storage conditions ^15^. To address this, we herein pair untargeted metabolomic-lipidomic profiling with functional assays to determine whether short-chain alcohols – specifically ethanol – can lessen metabolic uncoupling that emerges under supercooled conditions, making it possible to benefit from the improved retention of ATP reserves.

## RESULTS

### RBC Quality Indices Suggest Ethanol Attenuates Injury Exclusively Under Supercooled Storage

Hemolysis increased during supercooling, but 4% ethanol markedly attenuated this rise. (**Fig. 1A–B**). Ethanol significantly reduced day 21 and day 42 hemolysis levels at both −4 °C (day 21, p = 0.008; day 42, p = 0.004) and −8 C (day 21, p = 0.007; day 42, p = 0.0001), with the effect at −8 °C proving considerably greater than at −4 °C (ΔHemolysis with EtOH: −1.06% at −4 °C; −8.519% at −8 °C). Still, day 42 hemolysis levels in the ethanol-containing conditions at −8 °C and at −4 °C remain significantly above the 4 °C control indicating ethanol only partially corrects the injury that is specific to supercooled storage (−4 °C, p = 0.0355; −8 °C, p = 0.0002). Moreover, at 4 C itself, ethanol has no impact on hemolysis, suggesting its protective mechanism is specific to supercooled conditions.

**Figure 1.**
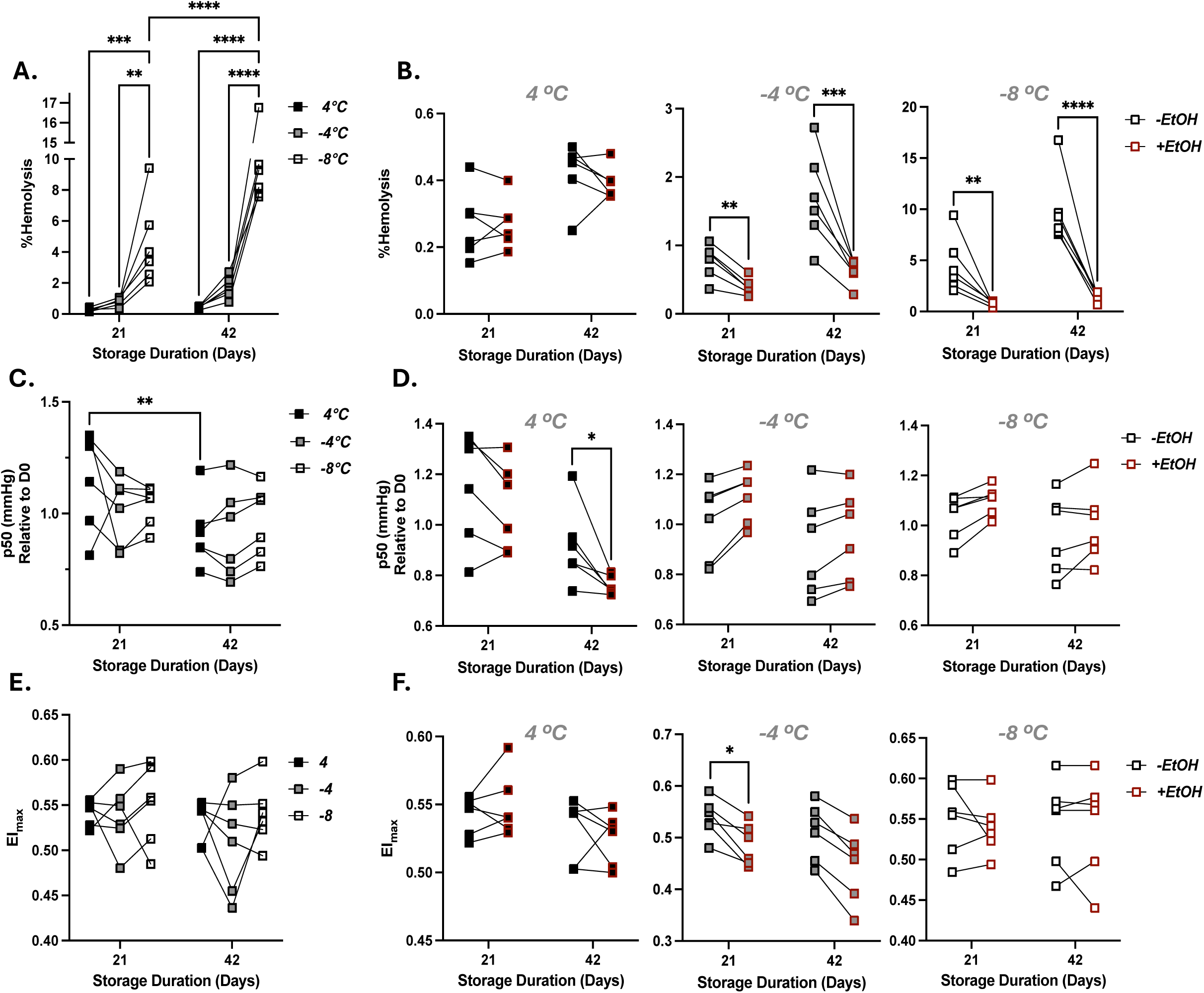
Ethanol modulates key quality metrics in red-blood-cells (RBCs) stored at 4 °C, –4 °C, and –8 °C. **(A, C, E)** Percent hemolysis, oxygen-carrying capacity, and deformability, respectively, for ethanol-free samples after 21 and 42 days. **(B, D, F)** The same parameters for ethanol-supplemented samples under the corresponding storage conditions. Statistics for panels A, C, and E, differences among temperatures at each time-point and temporal changes within a temperature were analysed by one-way ANOVA with Dunnett’s post-hoc test. In panels B, D, and F, ethanol-treated samples were compared with their paired ethanol-free controls at each temperature using two-tailed paired t-tests. In both cases: *p < 0.05, **p < 0.01, ***p < 0.001, ****p < 0.0001

Ethanol’s effects on other quality metrics were relatively modest (**Fig. 1C–F**). Across all storage temperatures, RBCs undergo a gradual morphological shift – from healthy discocytes to echinocytes and finally to spherocytes – a progression driven by microvesiculation and ionic or redox imbalances that disrupt membrane–cytoskeleton integrity (**Fig. S2**) ^17^. Ethanol does not alter morphology indices (a semi-quantitative score derived from subclass proportions) at any temperature or time point, yet at –4 °C, its presence significantly lowers the prevalence of discocyte subclasses: smooth discs (p = 0.0079), crenated discs (p = 0.049), and crenated discoid (p = 0.01). Consistent with the notion that discocyte-to spherocyte progression raises cytoplasmic viscosity and reduces membrane surface area, ethanol-treated samples at –4 °C exhibit a decrease in deformability (**Fig. 1F**) ^18^. These apparently adverse shifts (and any RBC quality comparisons between temperatures), however, must be interpreted with caution as the survivor bias can inflate the proportion of seemingly “healthy” cells.

Oxygen affinity, expressed as the P₅₀ ratio at day 21 (day 21) or day 42 (day 42) relative to baseline (D0), was not significantly altered by ethanol at either sub-zero temperature, although clear trends emerged (**Fig. 1C–D**; raw P₅₀ values in **Fig. S3**). At –4 °C and –8 °C, ethanol-treated samples showed higher P₅₀ ratios than controls at day 21, a trend that persisted to day 42 in five of six donors at –4 °C and three of six donors at –8 °C. By contrast, at 4 °C ethanol reduced P₅₀ ratios in three of six donors at day 21 and produced a significant reduction in all donors by day 42 (p = 0.0061).

### Ethanol Selectively Modulates Glycolysis and Purine Salvage Across Temperatures

Storage time and temperature dominated global metabolomic variance, with ethanol contributing modestly to group separation (**Fig. 2A; Table S1**). A pathway-oriented ROAST algorithm revealed subtler, physiologically relevant ethanol-driven shifts in glycolysis and purine salvage: pathways essential for RBC energy homeostasis (**Fig. 3A, B**).

**Figure 2.**
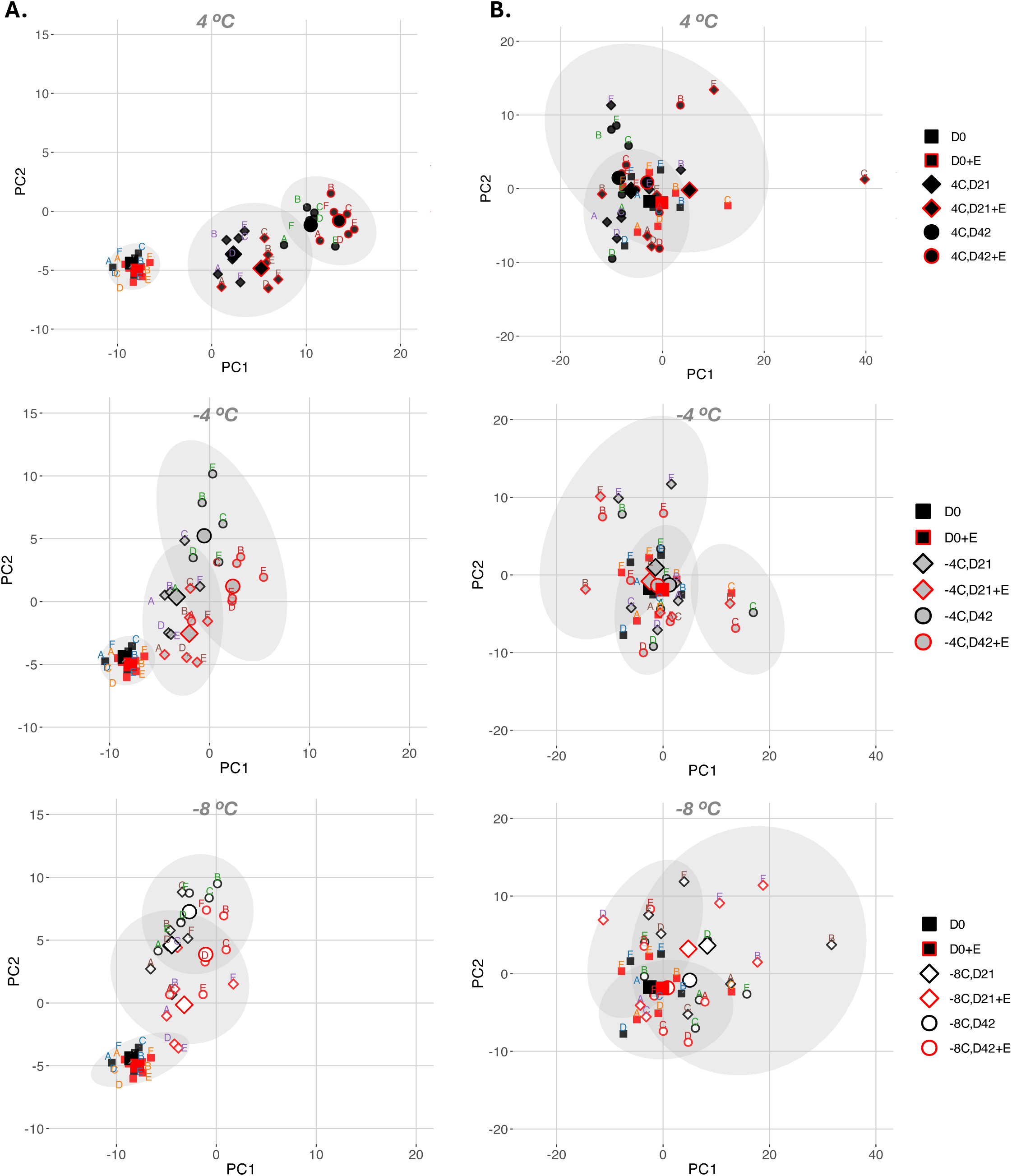
Multivariate analysis of metabolomic and lipidomic responses to ethanol and storage temperature**. (A)** Principal component analysis (PCA) with k-means clustering (k = 3) of the metabolomics data set. **(B)** Equivalent PCA-k-means plots for the lipidomics data set. Dashed black and red lines mark the median centroid distances of day-1 ethanol-free and ethanol-containing controls, respectively.

**Figure 3.**
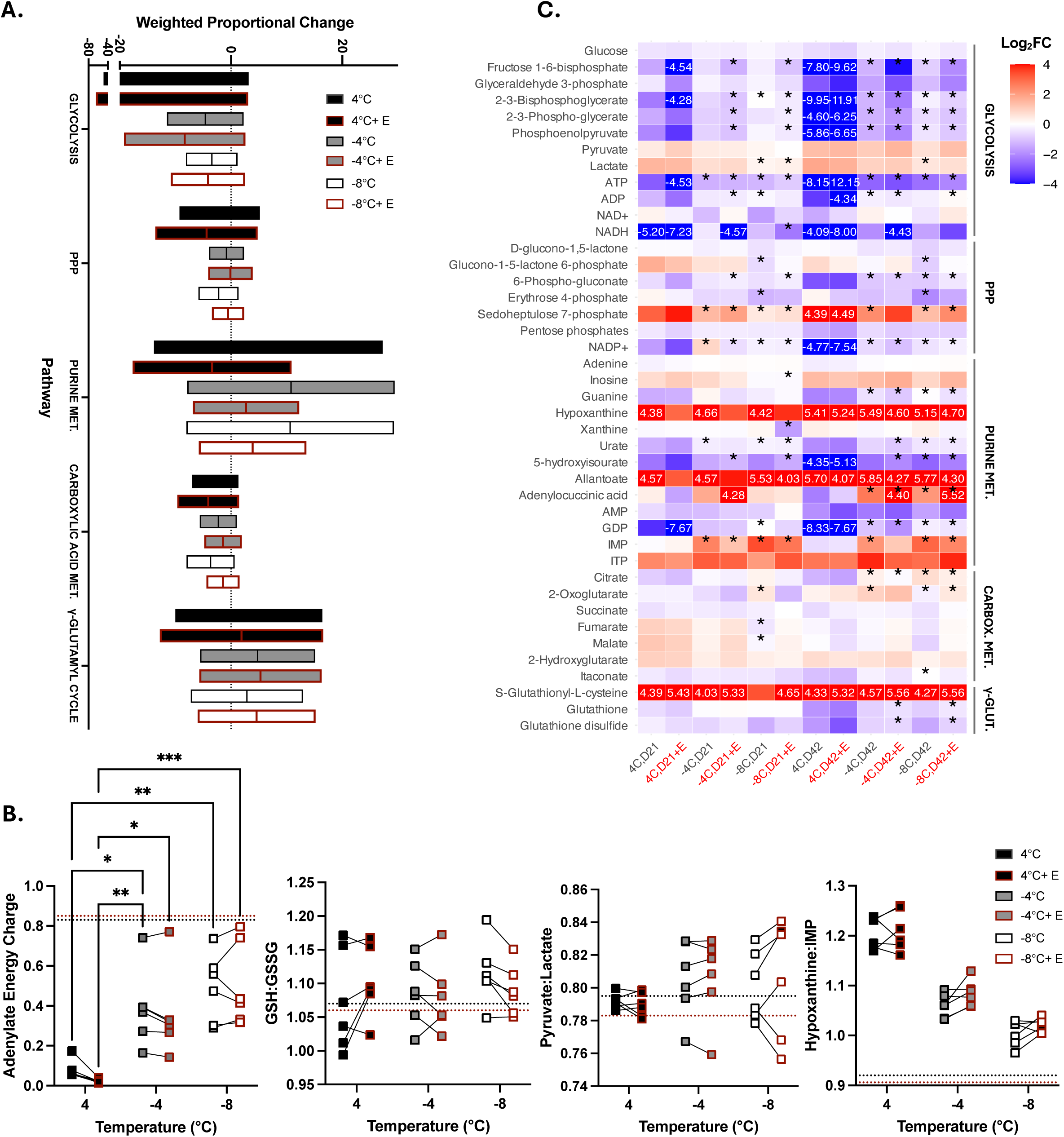
Temperature- and ethanol-dependent rewiring of RBC metabolism. **(A)** ROAST over-representation analysis summarizing the weighted proportional change (day 42 vs day 0) for each pathway; positive bars indicate net metabolite accumulation, negative bars indicate depletion, and line within the bars indicate the median. A similar plot with all analyzed metabolic pathways can be found in **Fig. S10**. **(B)** Metabolite ratios at day 42 reflecting adenylate energy charge (energy stored in the adenylate pool), hypoxanthine:IMP (purine salvage activation), pyruvate:lactate (late-stage glycolysis activation), and GSH:GSSG (cellular redox status). Significant differences across temperatures and ethanol treatments were calculated using as two-way ANOVA followed by a Dunnet’s post-hoc test: *p < 0.05, **p < 0.01, ***p < 0.001, ****p < 0.0001. **(C)** Heat map showing log₂-fold changes (log₂FC) for individual metabolites in the pathways outlined in panels A and B, calculated as log₂[(day 21 or day 42)/day 0].The colour scale is clipped at ±4 log₂FC; specific values beyond this range are displayed on the heatmap (heatmaps with all values included can be found in **Fig. S4**). For each metabolite, –4 °C and – 8 °C samples were compared with their 4 °C counterparts using two-tailed t-tests on log₁₀-transformed data (*p < 0.05).

Temperature and ethanol both strongly perturbed glycolysis (**Fig. 3A**). Results in **Figure 3** suggest that ethanol accelerates the energy-investment phase of glycolysis at 4 °C, with a similar but less pronounced effect observed at –4 °C and –8 °C. By day 42, significant reduction in 2- and 3-phosphoglycerate (p = 0.009), and phosphoenolpyruvate (PEP; p = 0.003) is apparent at 4 °C. Pyruvate levels and pyruvate to lactate ratios are, however, unaltered suggesting possible diversion of pyruvate into alternative pathways, such as those involving cytosolic isoforms of Krebs cycle enzymes (**Fig. 3A, C, S4**) ^19^. In keeping with the lack of increased flux through the energy payoff phase (i.e., PEP → pyruvate → lactate) despite the upstream acceleration of glycolytic flux, the adenylate energy charge (AEC) – a measure of metabolic energy stored in the adenylate pool – is significantly lower at day 21 in the ethanol-containing condition (**Fig. 3C; p = 0.022**). AEC at day 42 at 4 °C indicates ATP depletion irrespective of the presence of ethanol, but it is possible that ATP depleted sooner in the ethanol-containing condition (**Fig. 3C**). Conversely, at −4 °C and −8 °C, any changes in glycolytic flux in the presence of ethanol are ultimately inconsequential (**Fig. 3A**).

**Figure 4.**
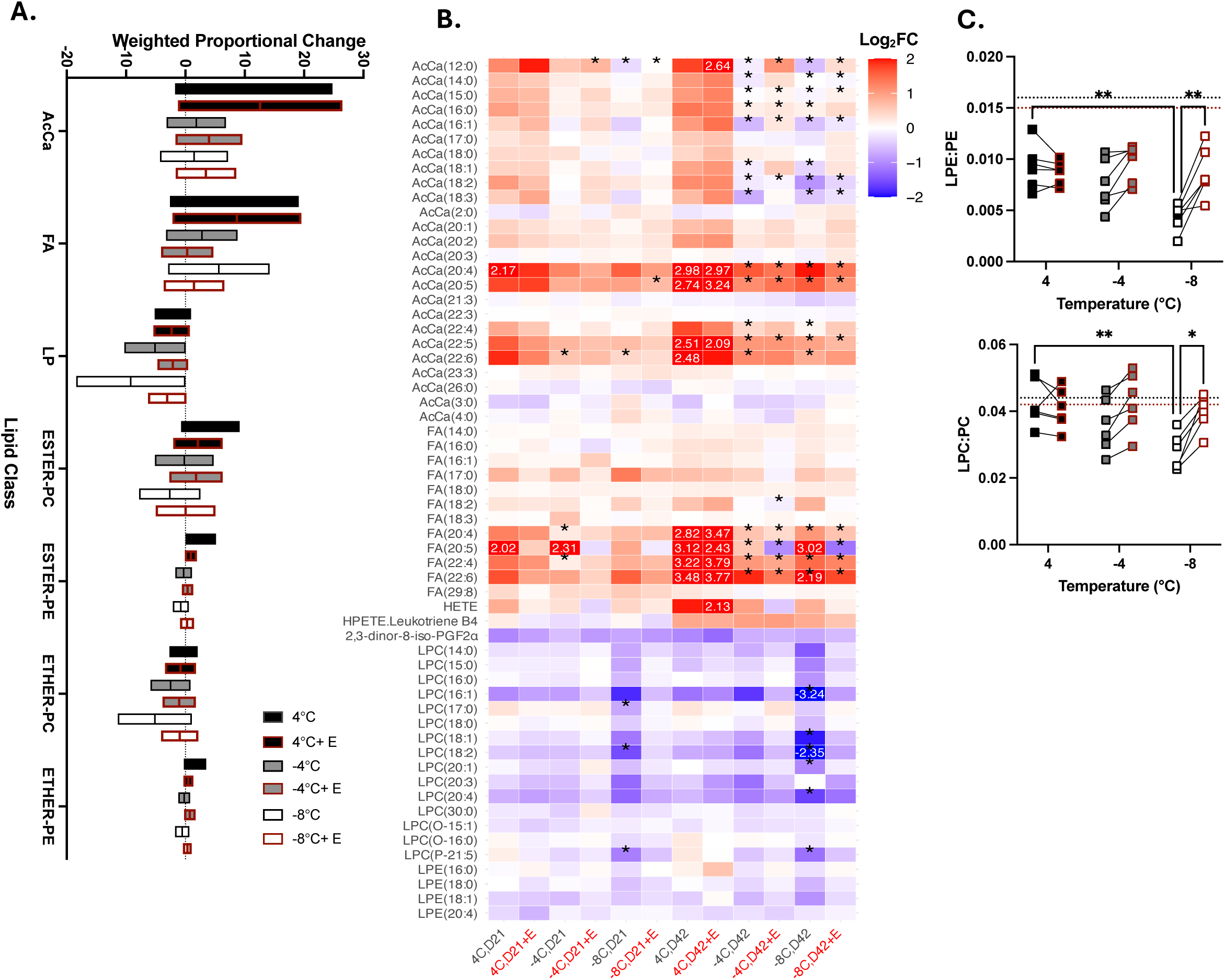
Temperature- and ethanol-dependent remodelling of the RBC lipidome. **(A)** ROAST pathway analysis showing the proportional day-42 change (vs day 0) for each pathway depicted in panel A. A similar plot with all analyzed metabolic pathways can be found in **Fig. S11**. **(B)** Heat maps of log₂FCs for classes of lipids experiencing the strongest changes across conditions based on panel (A), calculated as log₂(day 42/day 0). The colour scale is clipped at log₂FC ±4; specific values beyond this range are displayed on the heatmap (heatmaps with all values included can be found in **Fig. S4**). For each lipid, –4 °C and –8 °C samples were compared with their 4 °C counterparts using two-tailed t-tests on log₁₀-transformed data (*p < 0.05). Heat maps corresponding to all major lipid classes analyzed can be found in **Fig. S12**. **(C)** Ratios of total LPE:PE and LPC:PC at day 42. Symbols with red borders indicate the ethanol-supplemented condition. Significant differences across temperatures and ethanol treatments were calculated using as two-way ANOVA followed by a Dunnet’s post-hoc test: *p < 0.05, **p < 0.01. All data represents the mean of six biological replicates per condition.

While ethanol altered glycolysis, it had a negligible impact on the pentose phosphate pathway (PPP), purine salvage, and specialized nitrogen and sulfur metabolism. Despite temperature-driven shifts across these pathways (see Supplementary Notes 1 and 2), the steady-state abundance of key PPP intermediates and the hypoxanthine:IMP ratio (a reliable index of purine salvage flux) remained largely unaffected by ethanol (**Fig. 3A–C, S4**). A sharp, ethanol-induced decline in NADP⁺ at 4 °C effectively vanished under supercooled conditions (**Fig. 3A–C, S4**). Furthermore, while ethanol caused a slight (∼2–2.5 fold) increase in S-glutathionyl L-cysteine across all temperatures (**Fig. 3B, S4**), the canonical redox indicator GSH:GSSG remained stable at day 42, indicating that ethanol does not appreciably exacerbate oxidative stress (**Fig. 3C**).

### Ethanol Attenuates Lands’ Cycle-Driven Lipid Remodeling Exclusively Under Supercooled Conditions

Group separation driven by time, temperature, and ethanol in the lipidomics dataset was markedly smaller than in metabolomics (**Fig. 2B; Table S1**). Nevertheless, pathway analysis revealed supercooling-specific alterations in phospholipid remodeling that were attenuated by ethanol (**Fig. 4A–C**).

As RBCs are unable to synthesize glycerophospholipids through the Kennedy pathway, membrane phospholipid remodeling occurs primarily via the deacylation–reacylation cycle (i.e., the Lands’ cycle), mediated by phospholipase A2, PRDX6, and lysophosphatidyl acyltransferases ^20^. At 4 °C, ratios of lysophospholipids (LPs; deacylated products of phospholipids) to phospholipids – a proxy for Lands’ cycle activity – remain unchanged over 42 days of storage, irrespective of ethanol presence (**Fig. 4C**). Under supercooled conditions, however (particularly at –8 °C) these ratios are altered due to a decline in lysophosphatidylcholines (LPCs) and lysophosphatidylethanolamines (LPEs), a change largely attenuated by ethanol (**Fig. 4C**). Of the LP species that diminish during supercooling, all except LPE(20:4) and LPC(20:3) are higher when ethanol is present; for example, LPC(16:1) rises significantly by day 42 at –4 °C (p = 0.025), and LPC(16:1), LPC(18:1), LPC(14:0), and LPC(18:2) increase at –8 °C (all p < 0.0001) (**Fig. 4B**). ROAST analysis additionally indicates that ethanol partially offsets the accelerated loss of both ether- and ester-linked phosphatidylcholines and phosphatidylethanolamines during supercooled storage (**Fig. 4A**).

Ethanol similarly attenuated changes in select fatty acids (FAs) and acylcarnitines (AcCas) – lipid classes that contribute to the production of fatty acyl-CoAs for LP reacylation and that can accumulate when deacylation exceeds reacylation (**Fig. 4A, B**). Across all conditions (i.e., temperature, ethanol), long-chain polyunsaturated FAs (22:6, 22:4, 20:5, 20:4) and their corresponding AcCa species – several of which correlate strongly with post-transfusion recovery – exhibited the highest log₂ fold changes among all lipid species (**Fig. 4B**) ^21^. By day 42, levels of these species were highest at 4 °C, where ethanol had no appreciable impact (**Fig. 4B**). At day 42, these lipids peaked at 4 °C (where ethanol had no measurable effect) but were markedly lower at –4 °C and –8 °C (**Fig. 4B**). Moreover, ethanol further suppressed FA(22:6) and FA(20:5) under both supercooled conditions (p = 0.0004 for FA(22:6) at –4 °C; p < 0.0001 for all other comparisons), likely reflecting its inhibitory action on Δ6 and Δ5 fatty-acid desaturase (FADS2/FADS1) (**Fig. 4B**) ^22^. Ethanol also slightly, albeit insignificantly, reduced AcCa(20:5) and AcCa(20:4) levels at –4 °C and –8 °C, while minimally impacting AcCa(22:6) and AcCa(22:4) (**Fig. 4B**).

Lipid ontology (LION) enrichment analysis revealed that ethanol exerts a notable, temperature-dependent effect on the physical properties of lipids, including phase transition temperature (from liquid crystalline to gel), membrane charge, and curvature (**Fig. 5)**. At 4 °C, every class except “neutral headgroup” was negatively enriched in ethanol-treated samples; reaching significance for “very-high transition temperature,” “below-average transition temperature,” “very-high lateral diffusion,” “very-low transition temperature,” and “negatively charged headgroup.” At –4 °C and –8 °C, those same classes – apart from “very-low transition temperature” – showed a positive but non-significant enrichment. Notably, –4 °C samples displayed significant positive enrichment for “positively charged headgroup,” “high transition temperature,” and “above-average transition temperature,” whereas –8 °C samples showed significance only for “positively charged headgroup” and “positive intrinsic curvature.” The latter is of particular note as positive intrinsic curvature is a characteristic feature of lysophospholipids ^23^.

**Figure 5.**
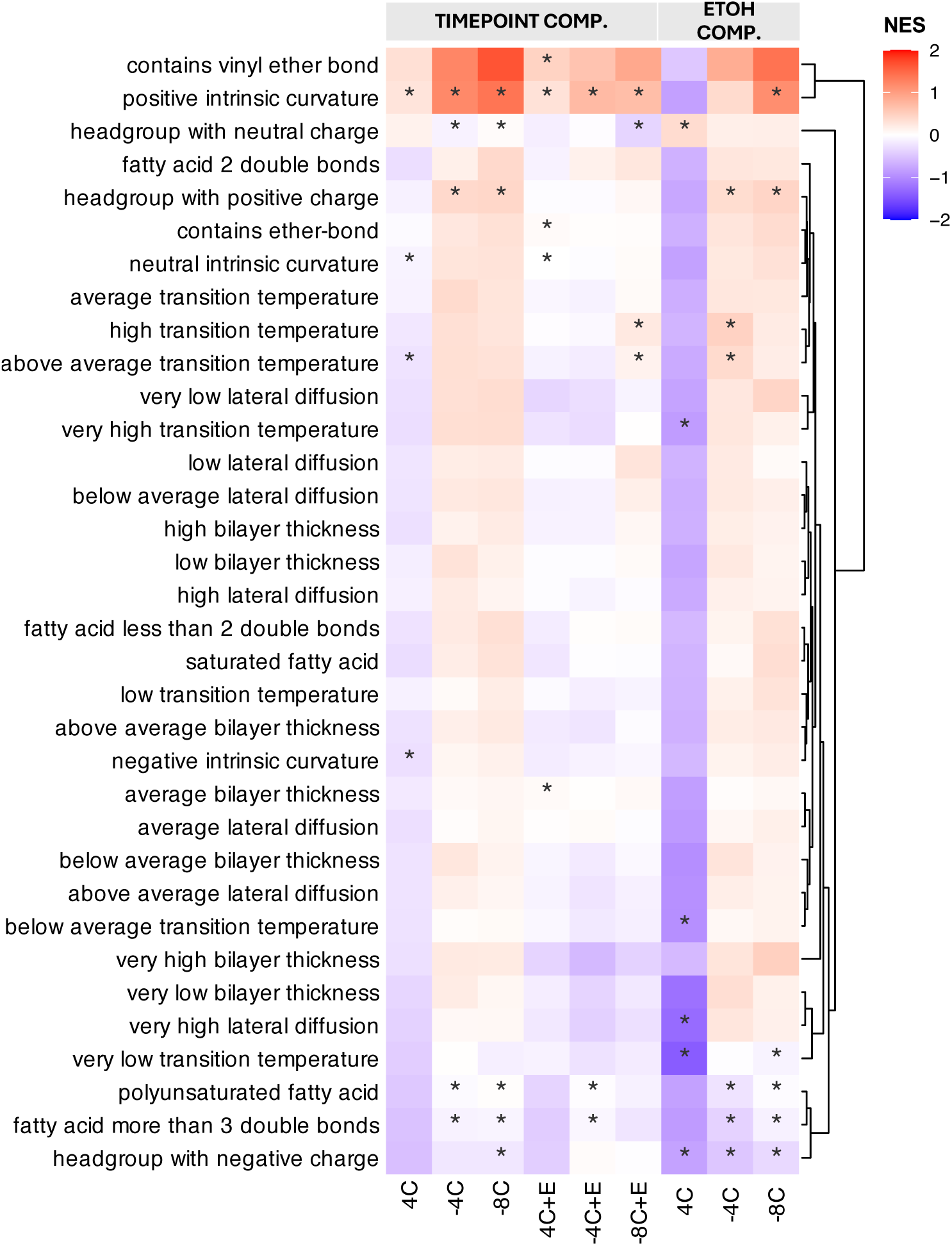
Lipid functional properties based on LION-ontology database. Normalized enrichment scores (NES) were calculated as the mean log₂-fold change (log₂FC) of all lipids within each term divided by the standard deviation across-lipids. Term significance relative to the background distribution was assessed with a two-sided Wilcoxon rank-sum test (*p < 0.05).

### Correlation Analyses Confirm Ethanol Reduces Lysophospholipid Loss at –8 °C and Exerts Broader Metabolic Modulation at –4 °C

Networks derived from weighted adjacency matrices – designed to suppress noise and capture coordinated monotonic trends across the integrated lipidomics-metabolomics dataset – exposed pathway-level shifts that were not apparent from individual metabolite fold-changes alone (**Fig. 6, Fig. S13**). Over-representation analysis of these matrices, restricted to pathways containing at least seven species to minimize instability and false positives, revealed pronounced ethanol-specific effects, most notably at –4 °C (**Fig. 6**). Here, enrichment scores – expressed as – log₁₀(p) from a hypergeometric test – indicated stronger coordination in branched-chain amino acid metabolism (isoleucine, leucine, valine), hydroxy-amino acid metabolism (glycine, serine, threonine), and glycolysis during ethanol exposure at –4 °C. Conversely, at –8 °C, ethanol exerted the opposite effect on alanine–aspartate–glutamate metabolism and reduced the enrichment of lysophospholipids (LPs), which aligns with the LP-specific log₂ fold-change analysis (**Fig. 4A, B**).

**Figure 6.**
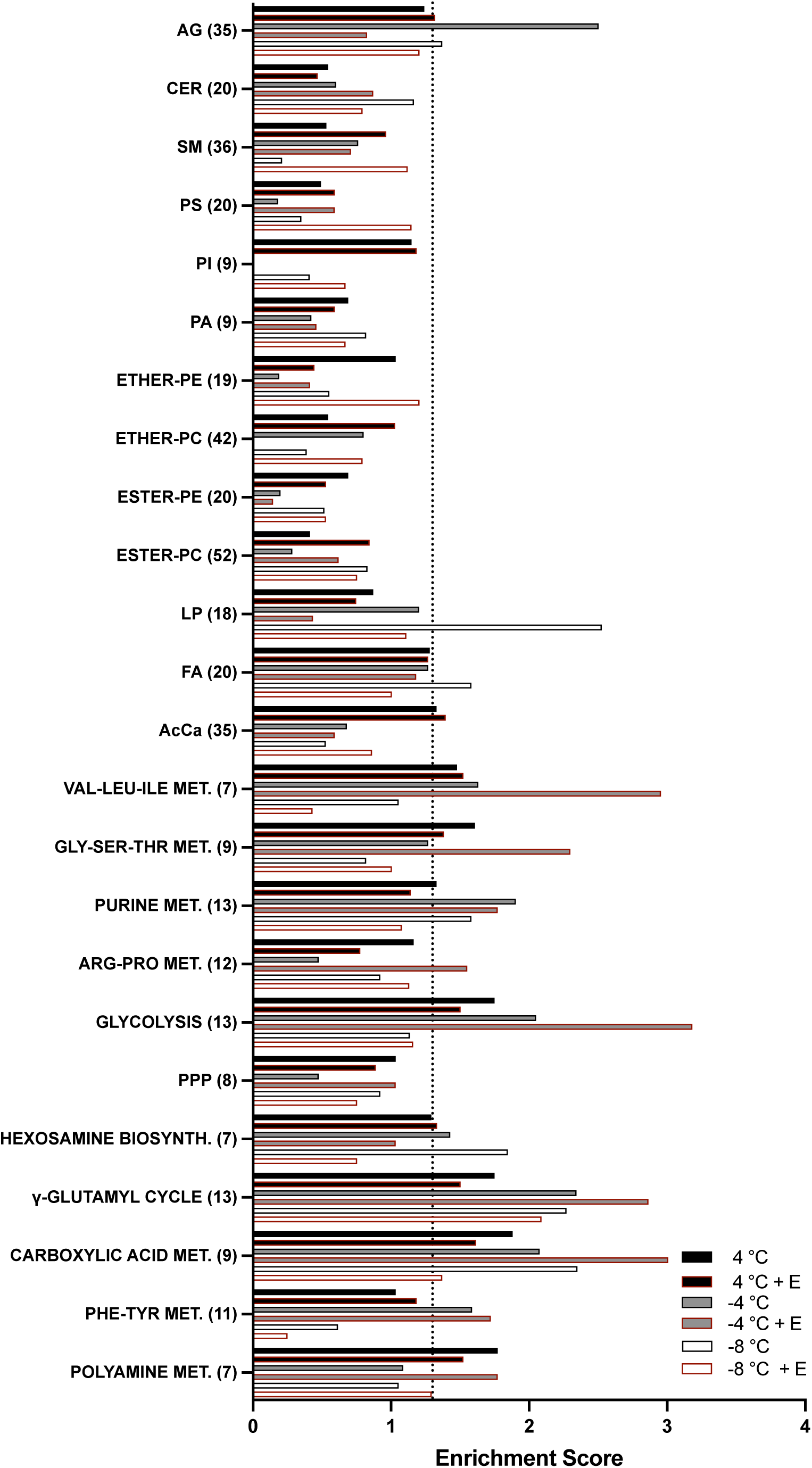
**P**athway and lipid class enrichment analysis of network-retained metabolites. Hypergeometric over-representation analysis (ORA) across storage temperatures (4 °C, –4 °C, and –8 °C) for both ethanol-free and ethanol-treated conditions, showing significantly enriched pathways and lipid classes for metabolites retained in the final adjacency matrices from Figure 6. The dashed line denotes the false discovery rate (FDR) significance threshold (-log_10_(FDR) > 1.3, equivalent to p < 0.05). Networks can be found in Fig. S13.

Rather than correlating metabolites and lipids with one another, correlating their shifts directly with hemolysis exposed temperature-dependent pathway vulnerabilities (**Fig. 7**). Over-representation analysis of these hemolysis correlates demonstrated that at 4 °C, hemolysis is strongly linked to perturbations in glycolysis, the γ-glutamyl-cycle, and AcCa’s (**Fig. 7B**). At –4 °C, the profile of hemolysis correlates expanded to include purine metabolism, fatty acids (FA), and lysophospholipids (LP), alongside continued significant enrichment in glycolysis. By –8 °C, however, this pattern simplified dramatically. The major metabolic pathways enriched at warmer temperatures were no longer significant; instead, lysophospholipids (LP) stood out as the sole, highly enriched class correlated with hemolysis at –8 °C (**Fig. 7B**).

**Figure 7.**
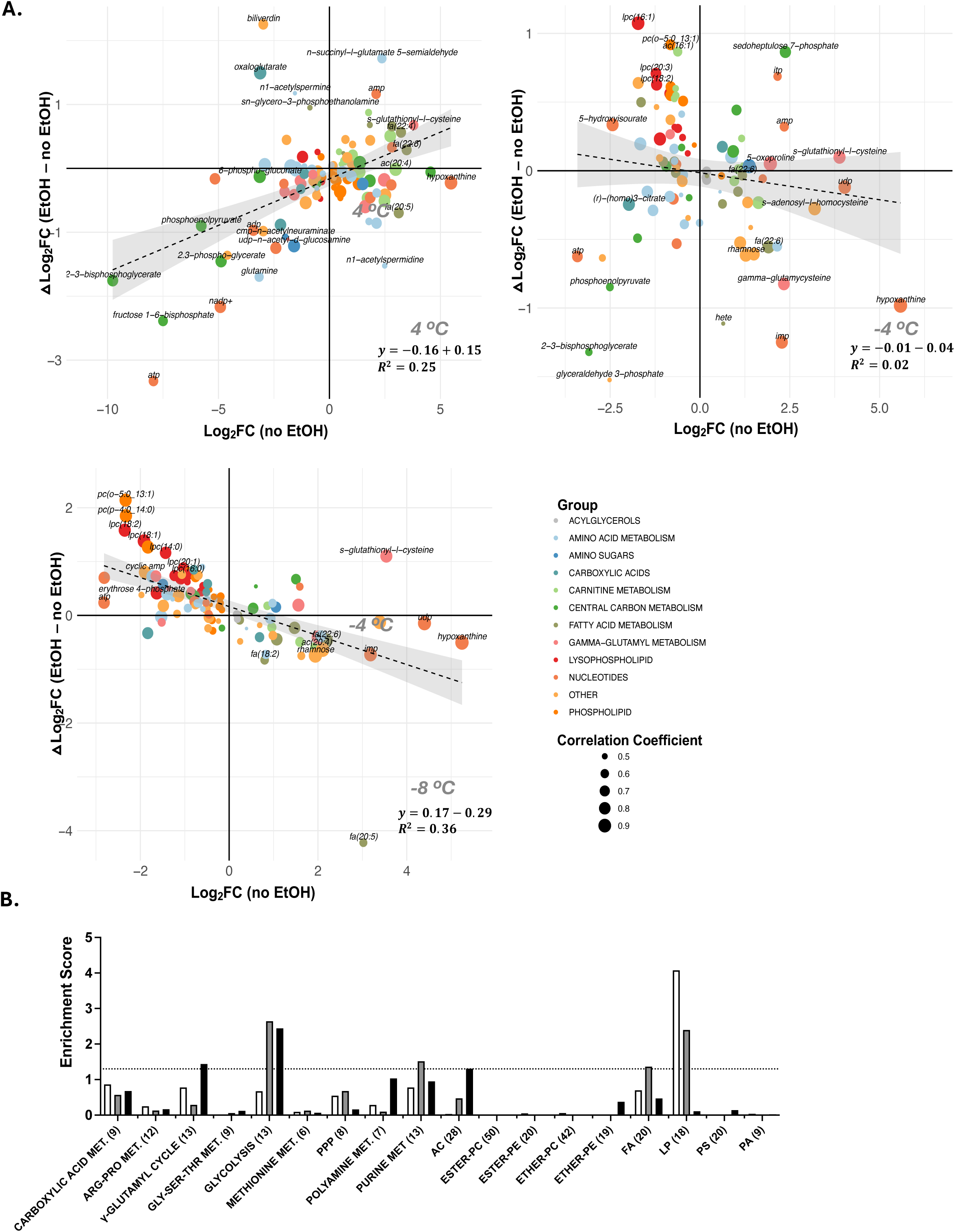
Ethanol mitigates hemolysis-linked metabolic and lipid perturbations as storage temperatures are lowered. (**A**) Scatter plots at day 42 for hemolysis-correlated metabolites across storage temperatures (4 °C, –4 °C, and –8 °C). The x-axis shows each metabolite’s log₂-fold change (log₂FC) relative to baseline (day 0) in the ethanol-free condition, while the y-axis shows the ethanol-driven shift (Δlog₂FC between ethanol-containing and ethanol-free conditions) at the same temperature and time-point. Solid lines are ordinary least-squares fits; shaded bands denote 95 % confidence intervals, with regression statistics inset in the lower-right of each panel. (**B**) Hypergeometric over-representation analysis of the metabolites in panel A, revealing pathways and lipid classes enriched among hemolysis correlates at each temperature. The dashed line indicates the +1.3 threshold that generally marks false-discovery-rate significance (p < 0.05).

## DISCUSSION

Supercooling preserves energy homeostasis during storage by attenuating catabolic processes; however, unlike during hypothermic storage, the resulting injury is not driven primarily by the loss of energy ^24^. Here, we identified that ethanol suppresses supercooling-specific injury in RBCs, markedly attenuating hemolysis at −4 °C and −8 °C while exerting a negligible effect at 4 °C. This protection occurred without major alterations to the overall metabolic signature, apart from a modest acceleration of glycolysis and reduced purine salvage (**Fig. 3**). Rather, the benefit of ethanol, particularly at −8 °C, stems from prevention of excessive loss of lysophospholipids and, to a lesser extent, phospholipids, which were strongly correlated with hemolysis under supercooled conditions (**Fig. 4**, **Fig. 7**). Because this supercooling-specific injury mechanism is previously unreported, we investigated its temperature dependence and ethanol’s mitigating role.

A loss of lysophospholipids may reflect activation of the Lands’ cycle of lipid remodeling, which aligns with homeoviscous adaptations (HVAs) where the membrane will remodel itself to increase fluidity and maintain function at low temperatures ^25,26^. This mechanism is common in poikilotherms; however, while mammalian cells undergo similar modifications, the homologous enzymes involved in HVAs lack equivalent low-temperature functionality ^27^. Thus, while the need for HVAs is theoretically more pronounced in a supercooled relative to a hypothermic state, the capacity of the cell to implement these modifications will be increasingly diminished. Several key aspects of our data support this interpretation. First, a reduction in a specific lysophospholipid would typically imply an increase in phospholipids with the same *sn*-1 chain, but this is not observed (**Fig. S12**). Second, there are no significant changes in the unsaturation indices of any phospholipid classes, which would be expected if the Lands’ cycle were enhancing low-temperature fluidity (**Fig. S5**). Moreover, lysophospholid degradation to produce fatty acids is unlikely to be driving its decline given that all fatty acid species decrease during supercooling, with no associated increase in AcCa species, (**Fig. 4A, B**). Instead, we posit that lysophospholipid loss is attributable to microvesiculation, a process which we know to increase in supercooled RBCs (**Fig. S6**) and can lead to hemolysis when excessive ^28^. At –8 °C, the increased retention of lipids with low transition temperatures and high lateral diffusion in the presence of ethanol further supports this idea, given that the more ordered membrane phases – comprising lipids with high transition temperatures and low lateral diffusion – are more prone to vesiculation.

There are both thermodynamic and biochemical factors conceivably driving heightened vesiculation under supercooled conditions, each of which ethanol could temper. Thermodynamically, membranes in the mixed-phase region of the thermotropic phase transition – where liquid crystalline (Lα) and gel (Lβ) phases coexist – experience line tension at their boundaries due to hydrophobic mismatch, creating an energetic penalty that can initiate microvesiculation ^29–31^. In RBCs, the main phase transition begins near 18 ± 2 °C, but the fluid Lα phase persists down to −10 to −5 °C because the membrane’s high cholesterol content (∼40 mol %) broadens the transition ^30,31^. Consequently, at −4 °C and −8 °C the membrane still contains both Lα and Lβ domains, and the line tension at their boundaries should exceed that at 4 °C. Lysophospholipids, with their conical shape, impose positive curvature at these boundaries, thereby reducing line tension but ultimately driving the vesiculation of lysophospholipid-rich membrane material ^23,32^. Although studies directly examining LP-induced attenuation of line tension between the Lβ and Lα phases are lacking, it is noteworthy that PLA₂ activity peaks at the onset of the thermotropic phase transition ^33^. If a heightened line tension does indeed drive vesiculation and LP loss under supercooled conditions, two mechanisms could explain ethanol’s protection. First, ethanol promotes the separation of headgroups and acyl chains, thereby reducing the phase transition temperature and decreasing the line tension that would otherwise develop between the Lβ and Lα regions at −4 °C and −8 °C ^34^. Second, the ethyl group of ethanol can interact with exposed acyl chain methyl groups at the domain boundaries, minimizing the thermodynamic forces that drive membrane budding and vesiculation ^35,36^.

At hypothermic temperatures, intracellular Ca²⁺ accumulation and oxidative stress are the chief biochemical drivers of vesiculation ^37^. Both damage the membrane and loosen its attachment to the cytoskeleton, thereby promoting vesiculation ^28,38^. Our preliminary data indicates an increase in calcium accumulation at –8 °C, and ethanol appears capable of attenuating this calcium buildup (**Fig. S7**). Because Ca²⁺-ATPase (the only channel facilitating Ca²⁺ extrusion in RBCs) is highly sensitive to local membrane order, its perturbation under supercooling conditions and subsequent activation by ethanol are readily explained ^39,40^. Additionally, ethanol may disrupt the autoinhibitory function of the C-terminal domain, thereby promoting Ca²⁺ release ^41^. The mechanisms by which ethanol might attenuate oxidative stress in RBCs are less well defined, especially considering that ethanol can be non-enzymatically oxidized to acetaldehyde, which in turn reacts with oxygen to generate hydroperoxyl radicals that promote lipid peroxidation ^42,43^. One could argue that by attenuating lateral phase separations, ethanol reduces the aggregation of unsaturated fatty acids and thereby slows the propagation of the peroxidation chain reaction; however, this hypothesis requires further investigation ^44,45^. Based on the metabolic signature (**Fig. 3**), it is challenging to determine whether ethanol is indeed alleviating oxidative stress, particularly as the metabolic signature of ethanol-free conditions suggest reduction in oxidative stress, which we know not to be the case from our prior studies showing attenuation of supercooling-specific injury using antioxidants (this decreased flux through pathways involved in oxidative stress tolerance instead likely reflects a reduced capacity to cope with oxidative stress) ^15^.

Regarding the biochemical and thermodynamic drivers of membrane loss, two possibilities exist for ethanol’s protective effect: (i) Does the elevated line tension outweigh the oxidative stress / calcium accumulation, making vesiculation an aberrant process rather than a protective strategy to clear damaged membrane material? and (ii) Might ethanol, by lessening line tension without adequately addressing the oxidative stress / calcium accumulation triggering vesiculation, allow damaged membrane material to accumulate excessively? This second point might suggest that a lower ethanol concentration than the 4% v/v used in this study would be more beneficial; however, data from **Fig. S8** indicate that concentrations of 1% and 2% are less effective at –8°C and –4°C in mitigating hemolysis. Interestingly, higher concentrations also prove beneficial, as 8% EtOH further reduces day 42 hemolysis at –4°C and –8°C (**Fig. S9**), prompting the question of where the upper limit lies at which the negative effects of ethanol addition (based on the abovementioned points) begin to outweigh its benefits. Ethanol, like all alcohols, increases membrane fluidity until a threshold is reached; beyond which, excessive headgroup separation creates energetically unfavorable voids that are mitigated by the interlocking of acyl chains from opposing leaflets, ultimately leading to increased membrane rigidity ^46^. In most cellular systems, this so called ‘ethanol-induced interdigitation’ typically occurs around 10–12% v/v, warranting further studies to determine the upper concentration limits and whether they depend on interdigitation ^47^.

This study advances high sub-zero (non-frozen) RBC storage as a complement to standard 1–6 °C storage, with potential to extend viable holding times for transport, pre-positioning, and surge capacity during disruptions. Mechanistically, we identify a lipidome-specific injury that emerges under supercooled conditions and show that low-dose ethanol can mitigate this membrane-centered damage. Extending this strategy to nucleated cells will require balancing membrane protection against additional supercooling stressors and alcohol-related toxicity: unlike RBCs, nucleated cells can oxidize ethanol to acetaldehyde, a reactive metabolite that forms adducts with nucleophilic groups and can promote radical chemistry ^48^. Secondary alcohols (e.g., 2-propanol, 2-butanol), which are oxidized to ketones rather than acetaldehyde, may therefore be better candidates for translating alcohol-mediated protection to nucleated cells and further optimizing RBC supercooling.

## MATERIALS AND METHODS

### Additive Solution Preparation

E-Sol 5 was used for optimal RBC supercooling ^15^. Prepared in-house, it contained 20 mM Na2HPO4, 25 mM citrate, 2 mM adenine, 111 mM glucose, and 41 mM mannitol (pH 8.4 at 37°C; ∼301 mOsm/kg). Ethanol (EtOH) was added at 1%, 2%, or 4% v/v on processing day. Solutions were vacuum-filtered (0.22 µm polyethersulfone membrane) prior to use.

### Blood Collection and Processing

Leukoreduced RBC concentrates (Canadian Blood Services) were washed three times within 3 days of collection to replace CPD-SAGM with EtOH-supplemented E-Sol 5, with the 4% v/v EtOH concentration confirmed to increase RBC membrane fluidity (Fig. S1). Concentrations were adjusted to 4.75 × 10^12 cells/L (45–50% hematocrit). RBCs were aliquoted (1 mL) into 5 mL tubes and sealed with 0.5 mL heavy paraffin oil to prevent nucleation. Following an overnight 4°C hold, supercooled samples were transferred to monitored portable freezers at −4°C or −8°C, with assessments occurring on days 0, 21, and 42. In vitro RBC quality parameters were evaluated using previously published methods. Specifically, hemolysis was measured optically via Drabkin’s method (Eq. 1.1), and hematological indices (MCV, MCH, and MCHC) were determined via a DxH 520 analyzer. Deformability (EImax) was derived using a Lorca analyzer at 37°C, while oxygen affinity (p50) was calculated via a Hemox analyzer for the 0% and 4% EtOH conditions only ^49^. Morphology was assessed via imaging flow cytometry (ImageStreamX Mark II), using a deep-learning classifier to categorize 5,000 images per sample into six subclasses to compute a morphology index ^50^. Finally, all data were analyzed via two-way ANOVA and Tukey’s post-hoc tests in GraphPad Prism (v. 10.4.0). condition.

### High-Throughput Metabolomics and Lipidomics

Supernatant and packed RBCs (0% and 4% EtOH; days 0, 21, 42) were frozen at −80°C. Aliquots (10 µL) were extracted in 96-well plates using 90 µL of ice-cold 5:3:2 MeOH:MeCN:water. After 0.2 µm filtration, extracts were analyzed via UHPLC-MS (Vanquish UHPLC coupled to a Q Exactive MS; Thermo Fisher) as previously described ^51^. Metabolites were resolved on a Phenomenex Kinetex C18 column at 45°C using a 1-minute ballistic gradient in positive and negative modes (65-975 m/z). The Q Exactive MS operated in negative Full MS mode (90–900 m/z, 70,000 resolution). Standardized blanks and quality controls were utilized to monitor instrument performance.

### Omics Data Analysis

Lipidomics and metabolomics data were combined and z-score normalized. Directional pathway enrichment was assessed via Rotation Gene Set Testing (ROAST) with proportion-based weighting. Lipid functional properties were analyzed using the LION-ontology database and Wilcoxon rank-sum tests. Key regulatory metabolites were identified by constructing weighted adjacency networks from bootstrapped Spearman correlations, optimized via gap statistic filtering, with eigenvector centrality calculated in Cytoscape. Hemolysis correlates were identified using Spearman analysis and hypergeometric overrepresentation. Analyses utilized R, MetaboAnalyst 6.0, and GraphPad Prism. Comprehensive mathematical and parametric details are provided in the Supplemental Methods.

## Institutional Review Board Statement

The collection and use of human blood samples were approved by the relevant research ethics boards at Canadian Blood Services and the University of Alberta (REB # 00095252 and REB #2021.013, respectively).

## CRediT

N.W.: Methodology, Investigation, Formal analysis, Data curation, Writing – original draft. Z.I.: Conceptualization, Writing – review & editing. T.N.: Data curation, Writing – review & editing. L.E.B.: Data curation, Writing – review & editing. M.H.: Data curation, Writing – review & editing. Y.Z.: Data curation, Writing – review & editing. J.K.: Data curation, Writing – review & editing. M.Y.: Data curation, Writing – review & editing. J.P.A.: Conceptualization, Funding acquisition, Project administration, Supervision, Writing – review & editing. A.D.: Project administration, Supervision, Writing – review & editing. O.B.U.: Conceptualization, Funding acquisition, Project administration, Supervision, Writing – review & editing.

