## Supplementary Figures for "Enhancing Supercooled Red Blood Cell Storage: The Membrane-Stabilizing Effect of Ethanol"

Table S1. Effect size measured based on partial eta squared values calculated from separate principal component analyses on metabolomics and lipidomics data.

| | | PC1 | PC2 | PC3 |
|---|---|---|---|---|
| **Metabolomics** | **Ethanol** | 0.02366433 | 0.07322435 | 0.00153484 |
| | **Time** | 0.39895686 | 0.34287385 | 0.10662694 |
| | **Temperature** | 0.13278903 | 0.49286673 | 0.02944814 |
| **Lipidomcis** | **Ethanol** | 0.00415103 | 0.00221962 | 0.01475404 |
| | **Time** | 0.01662014 | 0.02984517 | 0.02264452 |
| | **Temperature** | 0.10850842 | 0.01251376 | 0.00263881 |

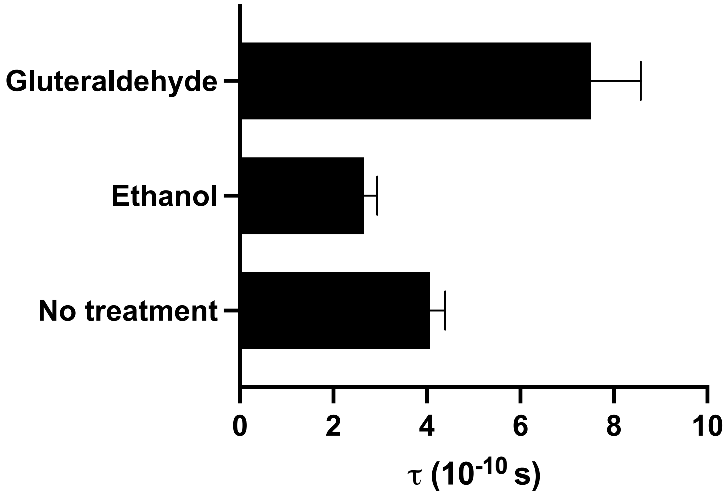


Figure S1. The effect of 4% ethanol on RBC membrane fluidity as determined by EPR spectroscopy evaluation of the rotational correlation time (τ) of 16-doxyl staeric acid. Untreated RBCs and RBCs treated with 0.5% (v/v) glutaraldehyde include for reference. Error bars in represent the standard deviation of the mean (n = 3).

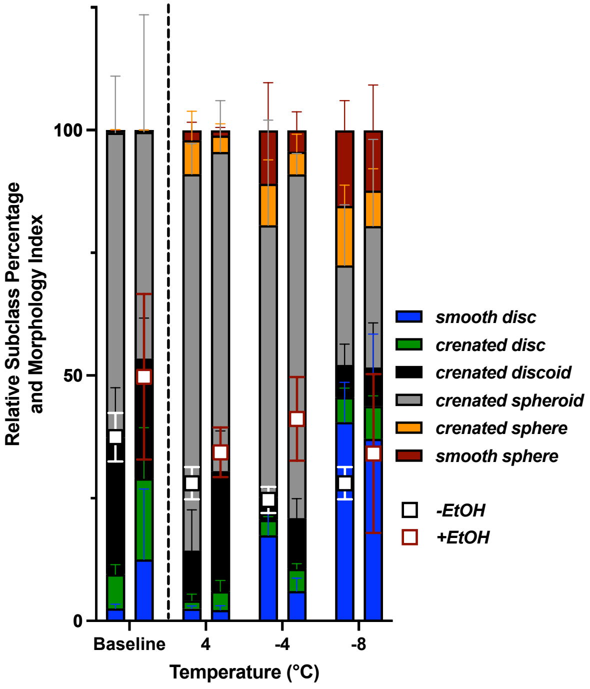


Figure S2. Morphological profile showing the proportion of each shape subclass and a composite morphology index calculated from weighted subclass fractions. Error bars  represent the standard error of the mean (n = 6).

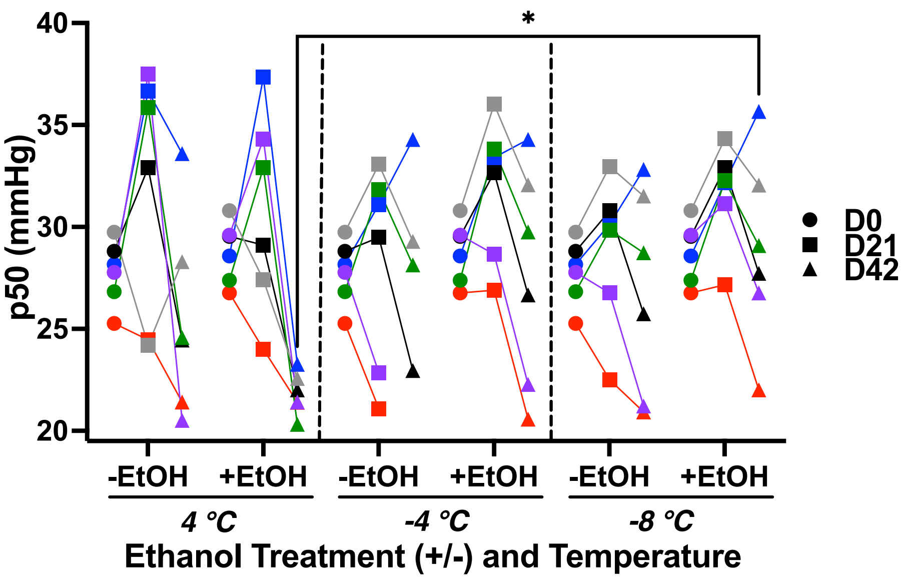


Figure S3. Impact of storage temperature and 4% ethanol on hemoglobin oxygen carrying capacity (based on partial pressure of oxygen: mmHg) measured after 21 and 42 days (labelled D21 and D42, respectively) of storage at 4°C, -4°C, and -8°C. The plot is color coded to indicate changes in individual samples across conditions.

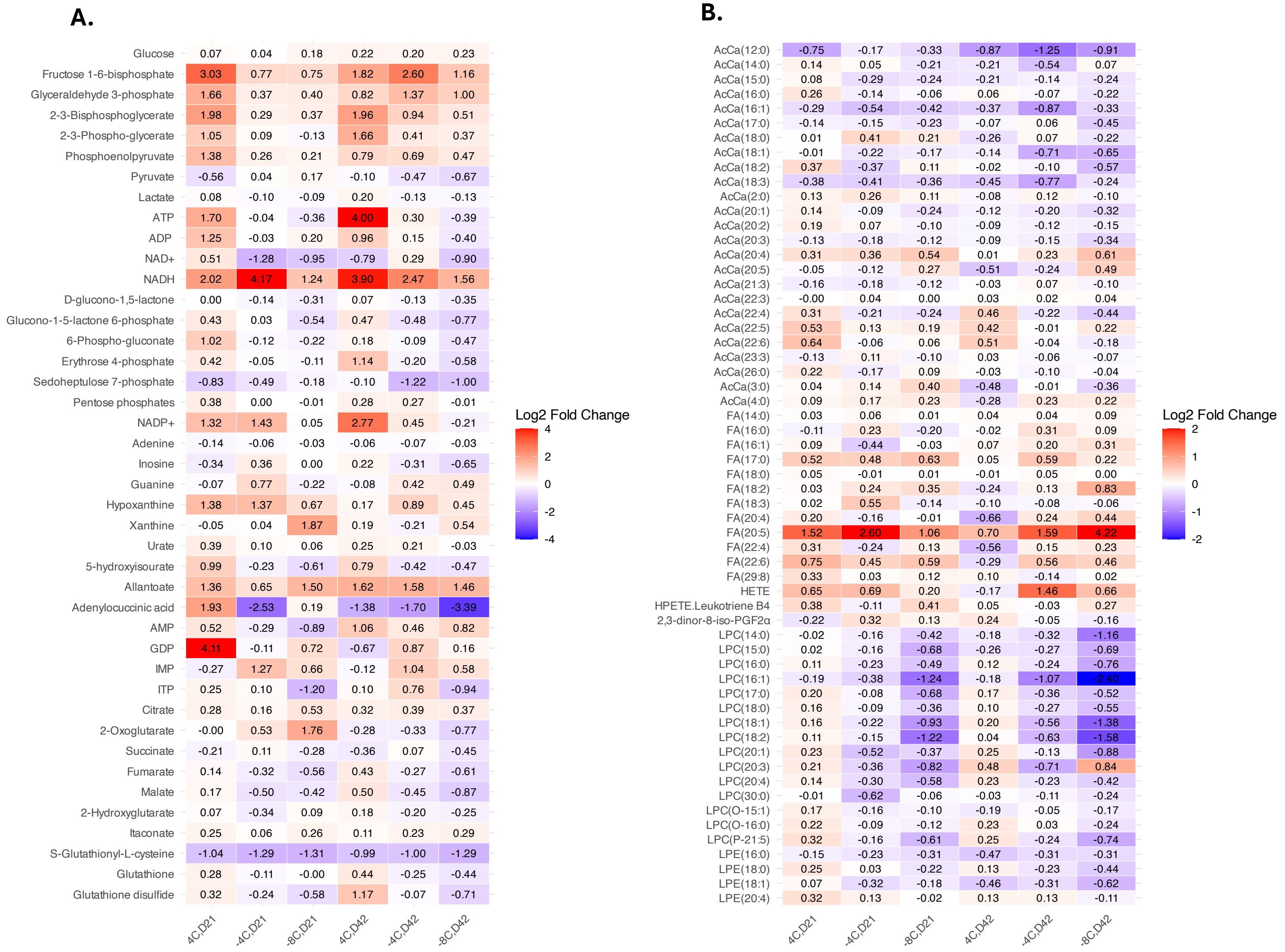


Figure S4. Impact of storage temperature and 4% ethanol on hemoglobin oxygen carrying capacity (based on partial pressure of oxygen: mmHg) measured after 21 and 42 days (labelled D21 and D42, respectively) of storage at 4°C, -4°C, and -8°C. The plot is color coded to indicate changes in individual samples across conditions.

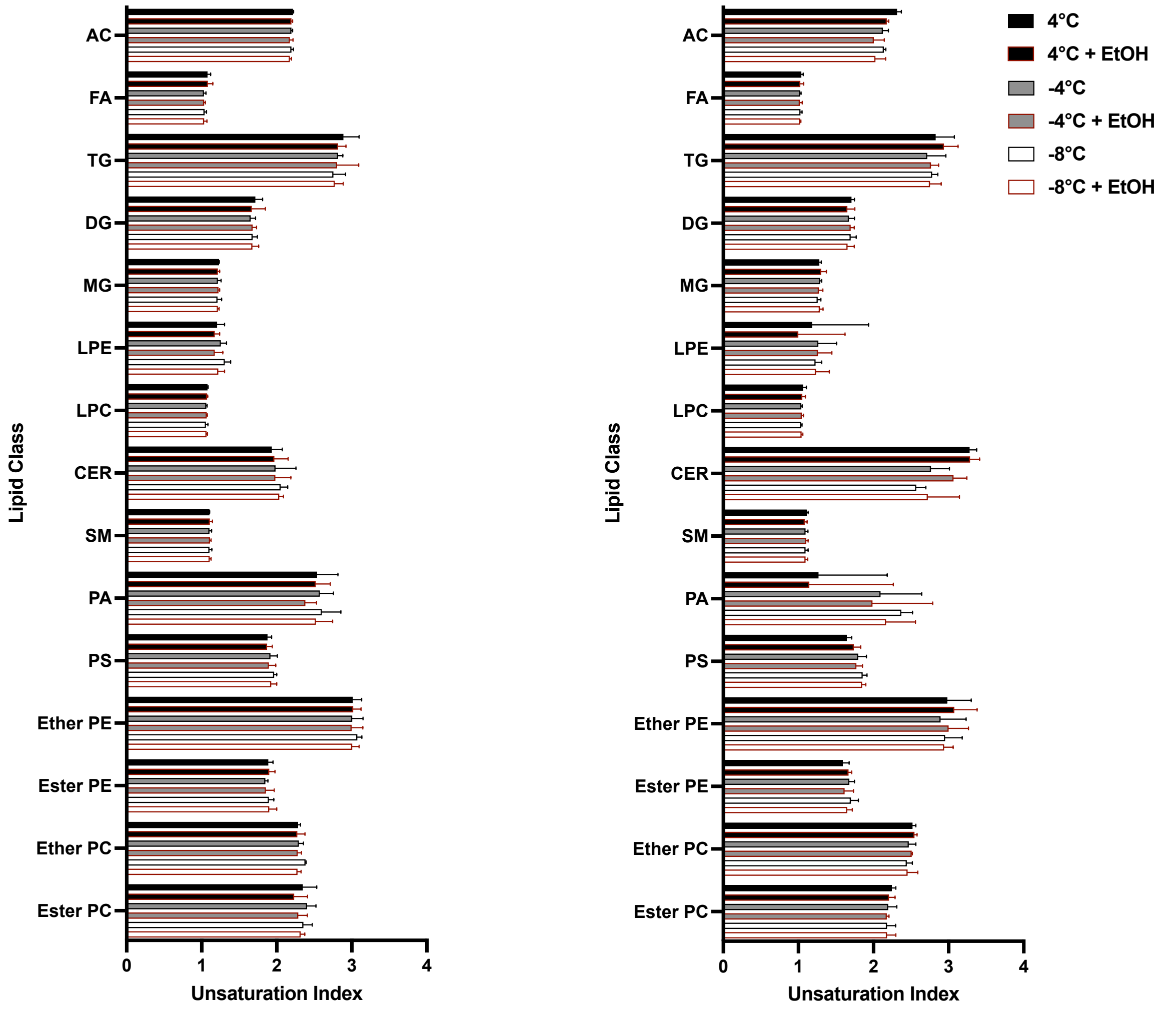


Figure S5. Unsaturation indices of lipid classes as determined using the following equation: UI = (sum of all lipids within class / weighted sum of all lipids within class)*100. The weighted sum is calculated by multiplying the quantified levels of each lipid species by the number of double bonds in the fatty acyl chains. Error bars in represent the standard deviation of the mean (n = 6).

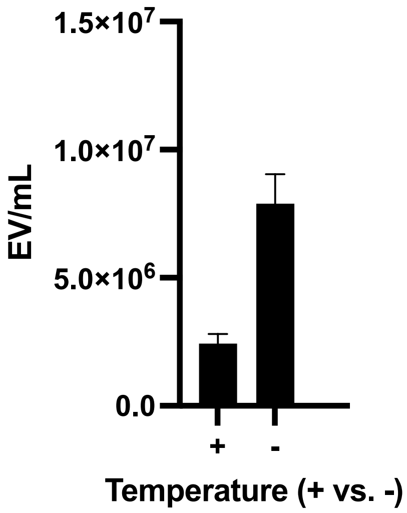


Figure S6. Extracellular vesicles in RBCs stored at 4C (+) and -8C (-) for 14 days. EV analysis performed through octadecyl rhodamine B chloride 18 staining and assessment through imaging flow cytometry.  Error bars indicate the standard deviation of three technical replicates from one biological replicate.

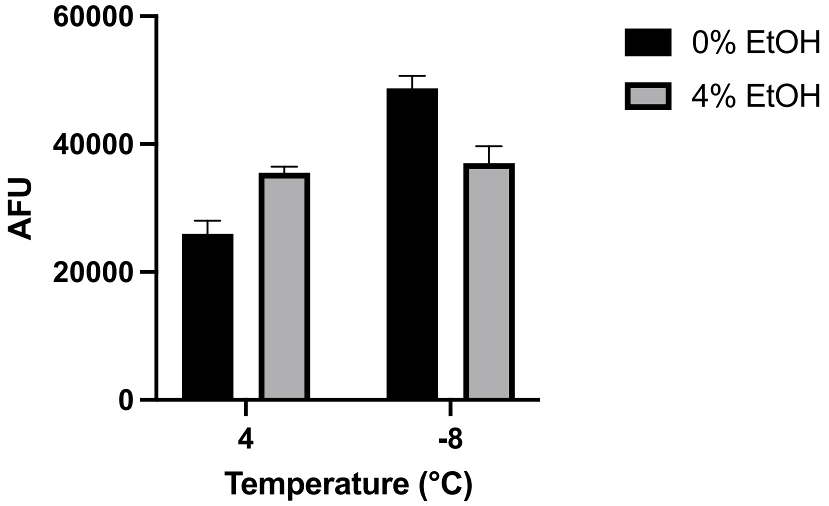


Figure S7. Calcium levels in RBCs stored for 42 days at 4 °C or -8 °C with 0% or 4% ethanol. Measurements were performed using imaging flow cytometry on RBCs stained with Cal500 AM. Error bars indicate the standard deviation of three technical replicates from one biological replicate.

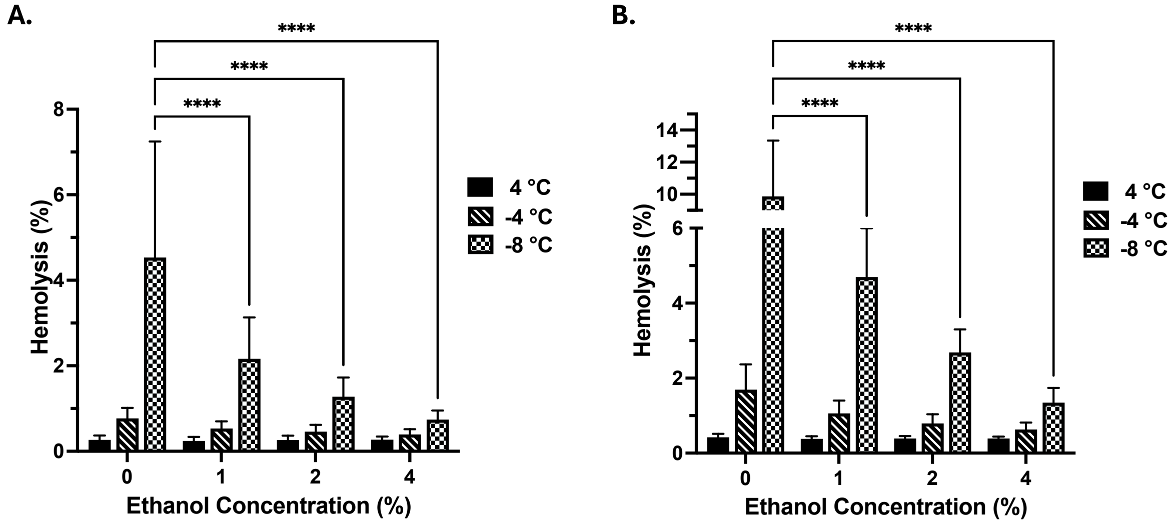


Figure S8. Impact of Low Ethanol Concentrations (1–4%) on Hemolysis During (A) 21-Day and (B) 42-Day Storage at 4 °C, -4 °C, and -8 °C. Significant differences calculated using a two-way analysis of variance (ANOVA) followed by a Tukey' post-hoc test to show differences relative to the 0% EtOH condition at each respective temperature: ****$p < 0.0001$. Error bars in represent the standard deviation of the mean ($n = 6$).

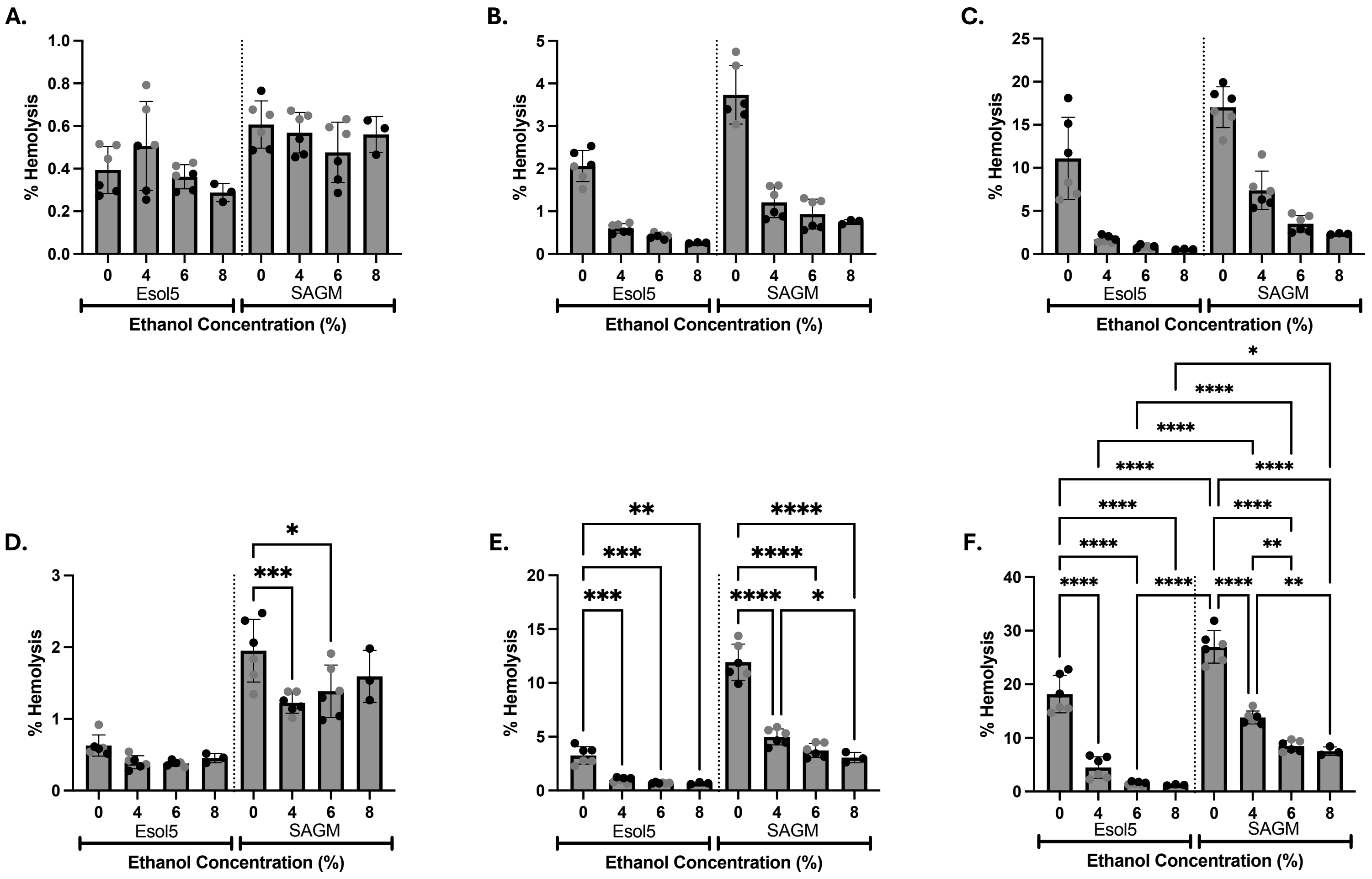


Figure S9. Impact of High Ethanol Concentrations (4–8%) and Storage Solutions (Esol5 vs. SAGM) on Hemolysis During 21-Day and 42-Day Storage at (A-B) 4 °C, (C-D) -4 °C, and (E-F) -8 °C. (A, C, E) Hemolysis for Esol5-stored RBCs; (B, D, F) Hemolysis for SAGM-stored RBCs. Error bars in represent the standard deviation of the mean (n = 6). Methods relevant to this figure are listed in the supplementary methods file.

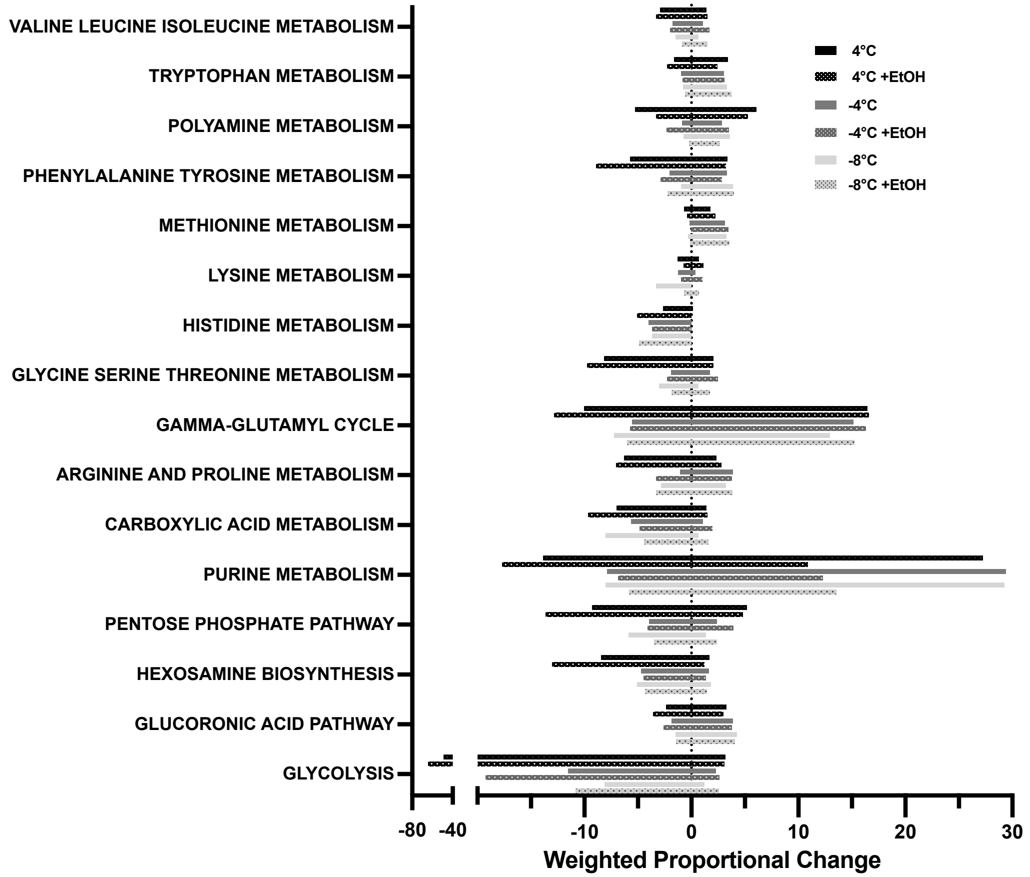


Figure S10. ROAST Analysis highlighting proportional changes in major metabolic pathways across 42 days of storage at 4 °C, -4 °C, and -8 °C. (n = 6)

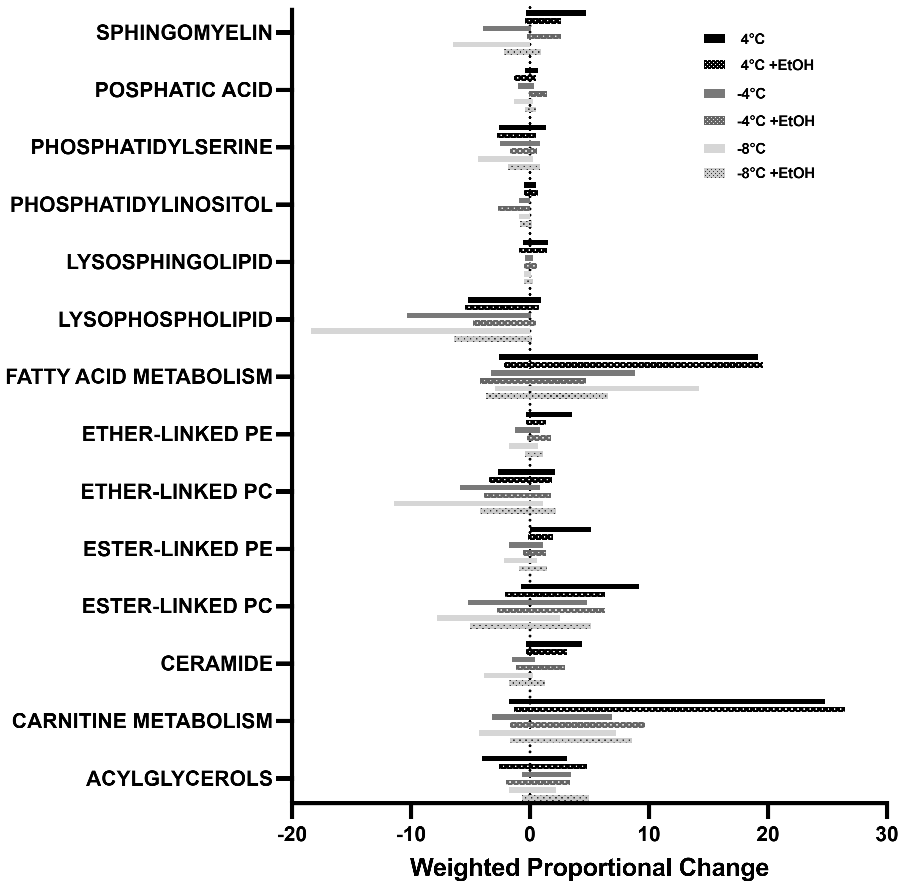


Figure S11. ROAST Analysis highlighting proportional changes in major lipid classes across 42 days of storage at 4 °C, -4 °C, and -8 °C. (n = 6)

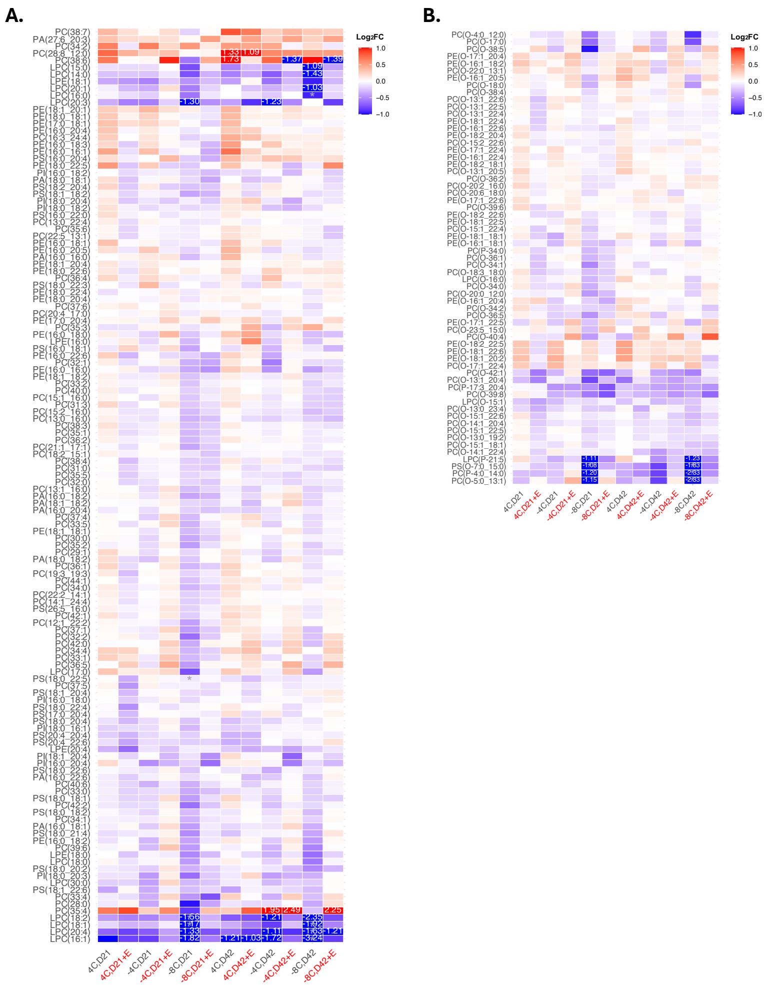

Figure S12. Heat maps of $\log_2$FCs for (A) ester lipids and (B) ether lipids, calculated as $\log_2$(day 42/day 0). The colour scale is clipped at $\log_2$FC ±1 for both panels; specific values beyond this range are displayed on the heatmap. For each lipid, –4 °C and –8 °C samples were compared with their 4 °C counterparts using two-tailed t-tests on $\log_{10}$-transformed data (*$p < 0.05$).

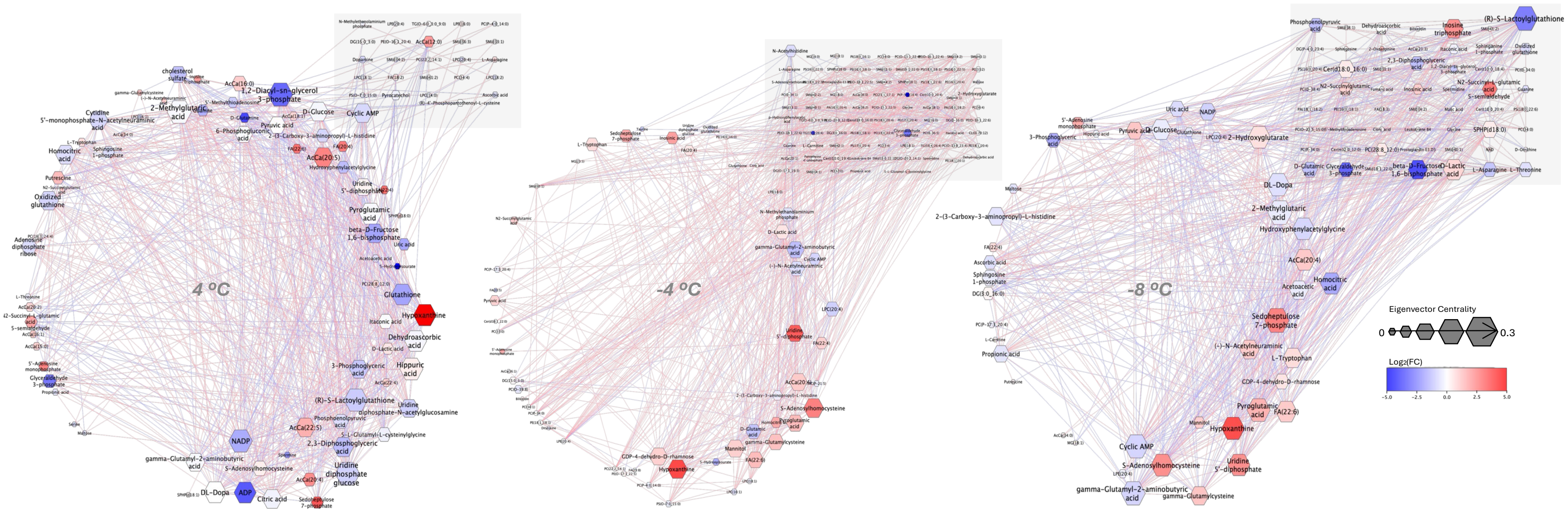


Figure S13. Network representations for ethanol-supplemented samples, corresponding to the ethanol-free networks shown in Figure 5. Each network retains the core topology of its counterpart; additional metabolites and lipids unique to the ethanol condition are boxed in gray (upper-right corner). The corresponding weighted adjacency matrix heatmaps for both figures are provided in Fig. S14.

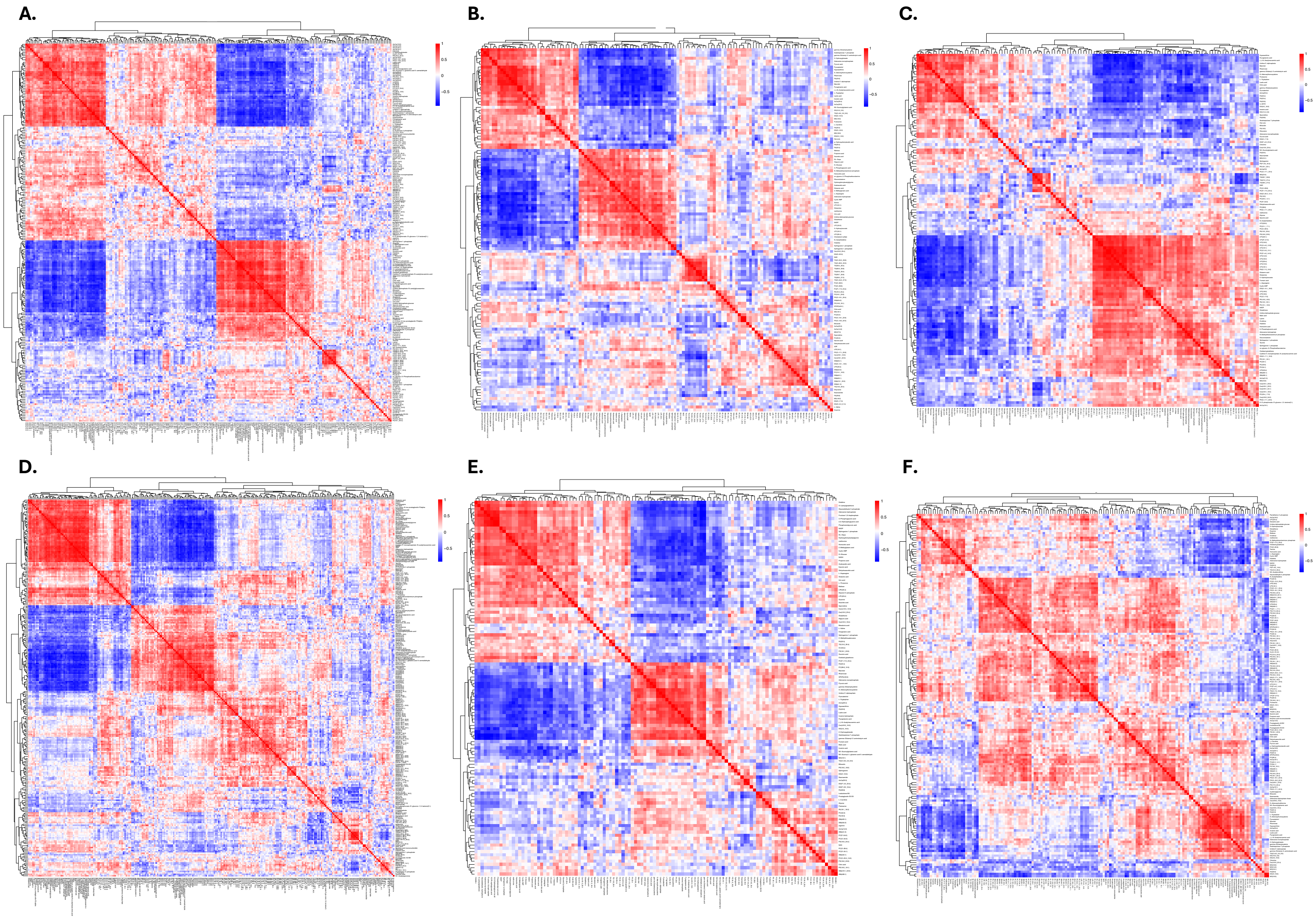


Figure S14. Heatmaps of weighted adjacency matrices constructed for all tested conditions following gap statistic filtration. Panels A–C show ethanol-free conditions at (A) 4 °C, (B) –4 °C, and (C) –8 °C, while panels D–F show ethanol-supplemented conditions at (D) 4 °C, (E) –4 °C, and (F) –8 °C.
