## Supplementary Methods for "Enhancing Supercooled Red Blood Cell Storage: The Membrane-Stabilizing Effect of Ethanol"

These notes offer a broader discussion of the methods used in the analysis of -omics datasets.

*Multivariate and Univariate Analyses*

Dimensionality reduction of the z-score normalized lipidomics and metabolomics datasets was performed using unsupervised principal component analysis (PCA) to evaluate within-group and between-group dispersion. To further explore meaningful between-group differences, k-means clustering (k = 3) was applied to the PCA-transformed data (2D space). Moreover, the effects of time, temperature, and ethanol treatments on separation along the first three principal components were determined through computation of partial eta squared squared values ($\eta^2$; a measure of overall effect size) as previously described [1]. In parallel, permutational multivariate analysis of dispersion (PERMDISP) quantified within-group variability to determine whether differences in dispersion influenced interpretations of between-group separation. PCA and both multivariate tests were conducted using "vegan" (R package; v. 2.6.8). Additionally, univariate analyses were performed on each -omics dataset to identify metabolites that differed significantly: (a) within each treatment group across the three tested temperatures (one-way ANOVA) and (b) between treatment groups at each temperature (paired t-test).

*Pathway-Level Enrichment Analyses*

To evaluate directional pathway-level enrichment across time points within each temperature/treatment condition, both datasets were combined, z-score normalized, and Rotation Gene Set Testing (ROAST) was performed using "limma" (R package; v 3.6.4) [2]. Unlike permutation tests, which disrupt the correlation structure of the null distribution and can obscure biologically meaningful co-regulated changes, ROAST applies random 'rotations' in multivariate space designed to preserve the correlational structure and interdependencies between metabolites (999 rotations were performed). The design matrix used in ROAST incorporated a pairing factor to account for samples derived from the same individual, ensuring that inter-individual variability did not introduce noise when computing net metabolite changes across time points. Eq. 1.1 indicates how the average abundance of a metabolite at each timepoint was calculated.

(1.1)
$$M_{i,Tx} = \frac{1}{N_x}\sum_{j=1}^{N_x} M_{i,Tx_j}$$

$M_{i,Tx}$ is the measured abundance of metabolite $i$ in sample $j$ at timepoint $x$ ($T_x$), where $x$ indicates timepoint 1, 2, or 3 (i.e., D0, day 21, or day 42). $N_x$ is the number of samples at $T_x$, which was constant across timepoints. From average abundance, Eq. 1.2 was used to calculate the net change in metabolites between timepoints.

(1.2)
$$\Delta M_i = (M_{i,T3} - M_{i,T1}) - (M_{i,T2} - M_{i,T1})$$

To prevent pathway size from biasing the results, a proportion-based approach was applied to metabolite up- and downregulation instead of absolute metabolite counts (see Eq. 2.3 and 2.4). When doing so, a weighting factor was introduced to refine the directional pathway enrichment scores, ensuring that each metabolite's contribution was proportional to the magnitude of its change across time points. For a given pathway $P$ containing metabolites $i$, the proportion of upregulated and downregulated metabolites was calculated as shown in Eq. 2.1 and 2.2, respectively.

(2.1)
$$\text{Proportion Up}_P = \frac{\sum_{i\in P} |\Delta M_i| \cdot I(\Delta M_i > 0)}{\sum_{i\in P} |\Delta M_i|}$$

(2.2)
$$\text{Proportion Down}_P = \frac{\sum_{i\in P} |\Delta M_i| \cdot I(\Delta M_i < 0)}{\sum_{i\in P} |\Delta M_i|}$$

$I(\Delta M_i > 0)$ is an indicator function that equals 1 if the metabolite increases across time, and 0 if it does not. Similarly, in Eq. 2.5, $I(\Delta M_i < 0)$ is an indicator function that equals 1 if the metabolite decreased over time, and 0 if it did not. $|\Delta M_i|$ represents the weighting factor, which is the absolute value of the metabolite's change.

*LION-Ontology Based Enrichment Analysis*

Detailed functional analysis of lipids was performed using classifications based on the LION lipid ontology reference database, which links lipids to three major branches: lipid class (based on LIPIDMAPS classification hierarchy), chemical and physical properties (i.e., fatty acid chain length / unsaturation, head group charge, and membrane fluidity / curvature effects), and major subcellular localization [3]. Only the terms within the chemical and physical properties branch were used, given the latter was irrelevant to RBCs and lipid class-based variation was already well-captured in prior analyses. A Normalized Enrichment Score (NES) was computed as the mean $\log_2$ fold change of lipids in the category, normalized by the standard deviation across all lipids. A Wilcoxon rank-sum test was then applied to assess whether LION group-associated lipids significantly differed from the background distribution.

*Network Construction and Centrality Analyses*

To identify key regulatory metabolites governing metabolic flux in each condition, network analysis was performed on aggregated, z-score normalized lipidomics and metabolomics data using a weighted adjacency matrix derived from bootstrapped Spearman correlations. This approach captured monotonic relationships over time while minimizing the impact of noise and outliers. 1,000 bootstrapped iterations were generated by resampling samples (donors/timepoints) while keeping metabolites / lipids fixed. Edge weights (metabolite-metabolite connections) in the final adjacency matrix were computed as the average correlation across bootstraps, divided by the frequency with which the edge appeared as a nonzero value across iterations.

To ensure biologically meaningful relationships were captured in the weighted adjacency matrix, a gap statistic approach was applied at two stages: (1) metabolite selection before bootstrapping, and (2) edge filtering after bootstrapping. SD thresholds were defined across 50th to 95th percentiles in 1% increments, with metabolites exceeding each threshold retained (e.g., at the 95th percentile, only metabolites with SD values higher than 95% of all metabolites were kept). At each quantile, a gap statistic was computed by subtracting the mean randomized variance (from 1,000 permutations) from the observed variance. These permutations disrupted correlation structure and metabolite interdependencies, generating a null distribution to estimate expected variance. The optimal SD threshold was identified at the so-called ‘elbow point’ of the gap statistic curve, ensuring only highly dynamic metabolites were retained for network construction.

After constructing the weighted adjacency matrix, a second gap statistic filtering step was applied to remove weak or unreliable edges. Here, absolute edge weights (averaged Spearman correlation values from 1,000 bootstraps) were used as filtering criteria. At each edge weight threshold, the total edge strength in the observed network was compared to that in randomized networks (generated via edge-weight shuffling). The gap statistic was computed as the difference between observed and mean randomized edge weights, allowing the identification of strong, biologically relevant metabolite-metabolite associations. As with metabolite selection criteria, the elbow point chosen as the cutoff, ensuring that only stable and reproducible edges were retained in the final network. The final weighted adjacency matrix was imported into Cytoscape (v3.10.2) for network visualization and analysis. Eigenvector centrality, which quantifies the influence of a node based on the number, strength, and importance of its connections, was computed using the cytoHubba (v0.1) plugin, with node size scaled to reflect eigenvector centrality.

The complete set of metabolites retained in the final weighted adjacency networks was also subjected to the hypergeometric over-representation procedure described below to providing pathway-level context for the global correlation network itself.

*Hemolysis Correlations and Overrepresentation Analyses*

To identify metabolite and lipid species significantly correlated with hemolysis at each temperature, Spearman correlation analysis was performed separately on z-score normalized data for samples stored at 4 °C, −4. °C, and −8 °C. Each analysis included day 0, day 21, and day 42 conditions, while ethanol-containing samples were excluded to better assess ethanol's contribution to significant correlates of hemolysis ($p < 0.05$). To determine which metabolic pathways or lipid classes were enriched among significant hemolysis correlates at each temperature, overrepresentation analysis was performed using a hypergeometric test. P-values were calculated using Eq. 3, where $K$ represents the number metabolites in the pathway, $n$ is the number of significant hemolysis correlates at each temperature, and $N$ is the total number of unique metabolites in the pathway reference file (Supplementary file 1) ($p < 0.05$ considered statistically significant).

(3)

$$p = \frac{\binom{K}{n}}{\binom{N}{n}}$$

*Software and Packages*

K means clustering and univariate statistical analyses were performed in Metaboanalyst 6.0, with all other analyses performed using RStudio (v. 2024-09-0+375). Graphpad Prism (v. 10.4.0) and "ggplot2" (R package; v. 3.5.1) were used to construct graphs. All heatmaps were contstructed using "pheatmap" (R package; v. 1.0.12). Other R packages used but not yet detailed, include: "dplyr" (v. 1.1.4), tidyr (v. 1.3.1), readr (v. 2.1.5), Hmisc (v. 5.2.2), stats (v. 4.4.1), factoextra (v. 1.0.7).


1. Pathmasiri, W., et al., *Integrating metabolomic signatures and psychosocial parameters in responsivity to an immersion treatment model for adolescent obesity.* Metabolomics, 2012. **8**: p. 1037-1051.
2. Liland, K.H., *Multivariate methods in metabolomics–from pre-processing to dimension reduction and statistical analysis.* TrAC Trends in Analytical Chemistry, 2011. **30**(6): p. 827-841.
3. Molenaar, M.R., et al., *LION/web: a web-based ontology enrichment tool for lipidomic data analysis.* Gigascience, 2019. **8**(6): p. giz061.
