## Supplementary Notes for "Enhancing Supercooled Red Blood Cell Storage: The Membrane-Stabilizing Effect of Ethanol"

These notes offer a broader discussion of metabolic pathway modulation with emphasis on the influence of temperature (rather than that of ethanol).

*SN-1. Modulation of Central Carbon Metabolism and Purine Salvage Pathways Under Supercooled Conditions*

A progressive decline in glycolytic capacity, despite adequate glucose availability, characterizes hypothermic storage injury in RBCs [1]. Temperature- and pH-induced impairments in glycolytic flux disrupt the NAD(H) redox balance and reduce ATP production, thereby disturbing cation and redox homeostasis and further compromising glycolysis [2]. These metabolic disturbances, along with the cells' diminished capacity to restore homeostasis under hypothermic conditions, result in decreased deformability and the development of spherocytic phenotypes [1]. Consistent with our recently published findings, lactate and pyruvate accumulate while upstream glycolytic intermediates are depleted across all storage conditions, with glycolysis emerging as the pathway most affected across temperatures (Fig. 3B and Fig. 3C) [3].

Supercooling markedly suppresses glycolysis, an effect more pronounced at –8 °C than at –4 °C (Fig. 3B and Fig. 3C). Although pyruvate and lactate levels decline slightly, the levels of glycolytic intermediates upstream of pyruvate—except for G3P—are significantly higher at –4 °C and –8 °C compared to 4 °C by D42 (Fig. 3C). Reduced flux through the energy-investment phase might compromise energy homeostasis by limiting substrates for the energy-payoff phase; however, depletion of ATP and ADP is less severe during supercooling. By day 42, the adenylate energy charge (AEC)—a measure of metabolic energy stored in the adenylate pool—is significantly higher at –4 °C and –8 °C than at 4 °C ($p = 0.0013$ and $p < 0.0001$, respectively), where an AEC of 0.1 reflects near-complete ATP depletion (Fig. S1). Moreover, supercooled storage preserves levels of 2,3-BPG, the positive allosteric regulator of Hb-$O_2$ affinity, suggesting a reduced reliance on its breakdown to support the energy-payoff phase (Fig. 3C).

Operating in parallel with glycolysis are the oxidative and non-oxidative phases of the pentose-phosphate pathway (PPP), which function respectively to restore redox homeostasis and supply substrates for the energy-investment phase of glycolysis (Fig. 3A). Diversion of glucose-6-phosphate into the oxidative phase generates NADPH, while diversion into the non-oxidative phase regenerates glycolytic hexose-monophosphate intermediates (glyceraldehyde-3-phosphate [G3P] and fructose-6-phosphate [F6P]) [50].

At 4 °C, oxidative PPP activation is evidenced by increased levels of glucono-1,5-lactone-6-phosphate (6PGL; $\log_2$FC, D42 = 0.78), the oxidation product of glucose-6-phosphate, accompanied by a decline in $NADP^+$ levels ($\log_2$FC, D42 = –4.76) (note that NADPH was not measured) (Fig. 3C). Although the levels of the end products of the oxidative PPP—the isobaric pentose-phosphate isomers (ribose, ribulose, and xylulose 5-phosphate)—decrease ($\log_2$FC = –1.46), this likely reflects their rapid utilization in other metabolic pathways. Across all temperatures the same pattern emerges, but oxidative PPP activation is generally lower under supercooled conditions. Specifically, 6PGL levels are significantly lower at –8 °C compared to 4 °C ($p = 0.049$), while $NADP^+$ levels are elevated at both –4 °C and –8 °C ($p < 0.0001$ in both cases).

In the non-oxidative phase of the PPP, transketolase catalyzes the conversion of ribose-5-phosphate (R5P, derived from the isomerization of ribulose-5-phosphate [Ru5P]) and xylulose-5-phosphate (X5P, formed via Ru5P epimerization) into G3P and sedoheptulose-7-phosphate (S7P) (Fig. 3A). These intermediates can be recycled to regenerate X5P and R5P or converted into erythrose–4-phosphate (E4P) and F6P via a reversible transaldolase-catalyzed reaction (Fig. 3A). Notably, S7P and E4P exhibit contrasting $\log_2$FCs at both D21 and D42: S7P levels consistently increase across all temperatures (D42 $\log_2$FC = 4 °C: 4.39; –4 °C: 2.34; –8 °C: 1.38), whereas E4P levels either decrease (at –4 °C and –8 °C) or remain effectively unchanged (at 4 °C) (D42 $\log_2$FC = 4 °C: 0.00005; –4 °C: –1.09; –8 °C: –2.03) (Fig. 3C). This pattern suggests that the reverse transaldolase-catalyzed reaction, which regenerates G3P and S7P, is favored over the forward reaction.

In addition to replenishing glycolytic intermediates and restoring redox balance, ribose-5-phosphate (R5P) produced by either branch of the PPP serves as the sole source of phosphoribosyl-pyrophosphate (PRPP), a critical substrate for purine-salvage pathways [4, 5]. This function is essential for maintaining nucleotide homeostasis in RBCs, which lack de novo synthesis pathways. Although changes in isobaric pentose phosphates do not suggest significant differences in purine salvage between temperatures, pathway-specific metabolites indicate that supercooling may promote purine salvage at the expense of catabolism.

In AMP catabolism, AMP is first deaminated to inosine monophosphate (IMP), which is then dephosphorylated and hydrolyzed to form hypoxanthine [6]. Because hypoxanthine can be rephosphorylated to IMP via PRPP, an increased IMP-to-hypoxanthine ratio—when AMP levels remain constant—indicates enhanced purine salvage. Notably, AMP levels are similar across temperatures, and while this is also the case for hypoxanthine, significantly higher IMP levels at –4 °C and –8 °C ($p < 0.0001$ in each case) result in elevated IMP-to-hypoxanthine ratios at both D21 (4 °C: 0.19; –4 °C: 0.96; –8 °C: 2.1) and D42 (4 °C: 0.06; –4 °C: 0.42; –8 °C: 0.97) (Fig. 3C). Given the cellular requirements for purine salvage would be diminished during supercooling due to the higher AEC, it is possible this seemingly counter-intuitive observation is the result of a bottleneck in the conversion of IMP to hypoxanthine.

*SN-2. Supercooling Suppresses Nitrogen and Sulfur Metabolism*

Red blood cells (RBCs) depend on several specialized metabolic pathways beyond central carbon metabolism to maintain redox balance and protein homeostasis, with nitrogen and sulfur metabolism playing crucial roles. As expected, supercooling suppresses these pathways—reducing catabolic flux and, in some cases, indicating a diminished ability to cope with stress (Fig. 3B).The methionine cycle repairs oxidized isoaspartate residues and, through its interplay with polyamine metabolism, enhances oxidative stress tolerance by scavenging reactive oxidants like hydroxyl radicals. S-adenosyl methionine (SAM), the primary methyl donor in the Protein-L-isoaspartate O-methyltransferase (PCMT1) system, remains stable across temperatures. However, its by-product, S-adenosyl homocysteine (SAH), increases under supercooled conditions (−4 °C and −8 °C). This rise may reflect impaired recycling of SAH to SAM, a diversion into the transsulfuration pathway, or enhanced PCMT1 activity in response to greater isoaspartyl damage.

Decarboxylated SAM—a derivative formed by removing SAM's carboxyl group—serves as a key aminopropyl donor that converts putrescine into spermidine and spermidine into spermine, with each reaction producing 5-methylthioadenosine (5-MTA) as a by-product (Fig. 3A). While decarboxylated SAM was not directly measured, 5-MTA levels generally decline under most conditions but remain significantly elevated by D42 at −4 °C and −8 °C (Fig. 3C). This pattern may suggest either increased activity of the methionine-salvage pathway at 4 °C or reduced polyamine synthesis under supercooling. At 4 °C, putrescine levels increase, while both spermidine and spermine decrease, with spermidine showing a more pronounced decline (Fig. 3C). In contrast, under supercooled conditions, spermidine levels remain stable, whereas putrescine is significantly lower and spermine is significantly higher (Fig. 3C). The concurrent rise in N[1]-acetylspermidine across temperatures, together with these changes, points to catabolism of polyamines (a process initiated primarily in response to polyamine oxidation); however, lower levels of N[1]-acetylspermidine at −4 °C and −8 °C suggest a reduction in polyamine catabolism under these conditions (Fig. 3C).

Mechanisms of nitrogen turnover offer additional insight into protein damage and amino-acid catabolism. A streamlined set of Krebs-cycle reactions among carboxylic acids supports nitrogen turnover while complementing the oxidative pentose-phosphate pathway (PPP) by providing an alternative route for NADPH regeneration under oxidative stress. Supercooling suppresses carboxylic-acid metabolism, as shown by the reduced breakdown of citrate and glutamine—two key contributors to alternative NADPH-generating pathways in RBCs. By Day 42, levels of both metabolites remain significantly higher at –4 °C and –8 °C compared with 4 °C ([citrate]: 4 °C vs –4 °C and 4 °C vs –8 °C, $p < 0.0001$; [glutamine]: 4 °C vs –4 °C, $p = 0.03$; 4 °C vs –8 °C, $p = 0.02$), indicating temperature-dependent suppression of their metabolism (Fig. 3C).

Reduced glutamate under supercooled conditions (the end product of glutaminolysis and the primary nitrogen donor in the cell) points to diminished nitrogen turnover that may stem from lower nitrogen assimilation or decreased amino-acid catabolism. Shifts in oxoglutarate, the chief nitrogen acceptor in transamination reactions, reinforce this conclusion; although its $\log_2$ fold change remains negative, the drop is less pronounced under supercooling (Fig. 3C). Arginine metabolism, which detoxifies and clears excess nitrogen, signals lower free-nitrogen levels at –4 °C than at –8 °C (Fig. 3C). Normally, limited nitrogen turnover would raise arginine because less nitrogen is routed into urea and creatine, yet our data show reduced arginine at –8 °C, while levels at –4 °C are similar to those at 4 °C (Fig. 3C). In line with this pattern, ornithine (produced by arginine hydrolysis) rises at –8 °C but not at –4 °C (Fig. 3C).
